# RNA processing machineries in Archaea: the 5’-3’ exoribonuclease aRNase J of the β-CASP family is engaged specifically with the helicase ASH-Ski2 and the 3’-5’ exoribonucleolytic RNA exosome machinery

**DOI:** 10.1101/699629

**Authors:** Duy Khanh Phung, Clarisse Etienne, Manon Batista, Petra Langendijk-Genevaux, Yann Moalic, Sébastien Laurent, Violette Morales, Mohamed Jebbar, Gwennaele Fichant, Marie Bouvier, Didier Flament, Béatrice Clouet-d’Orval

**Affiliations:** Laboratoire de Microbiologie et de Génétique Moléculaires, UMR5100, Centre de Biologie Intégrative (CBI), Université de Toulouse, CNRS, Université Paul Sabatier, F-31062 Toulouse, France; Université de Brest, CNRS, Ifremer, Laboratoire de Microbiologie des Environnements Extrêmes, F-29280 Plouzané, France

**Author notes:** These authors have participated equally to this work (Duy Khanh Phung, Clarisse Etienne). Corresponding Author orcid.org/0000-0002-3591-8538; Laboratoire de Microbiologie et Génétique Moléculaires-UMR 5100-Centre de Biologie Intégrative, CNRS-Université de Toulouse Paul Sabatier, Bât. IBCG, 118, route de Narbonne, 31062 Toulouse Cedex 9, France.

**Keywords:** Archaea, RNA metabolism, β-CASP Ribonuclease, Ski2-like helicase, ribosome, RNA exosome

## Abstract

A network of RNA helicases, endoribonucleases, and exoribonucleases regulates the quantity and quality of cellular RNAs. To date, mechanistic studies focused on bacterial and eukaryal systems due to the challenge of identifying the main drivers of RNA decay and processing in Archaea. Here, our data support that aRNase J, a 5’-3’ exoribonuclease of the β-CASP family conserved in Euryarchaea, engages specifically with a Ski2-like helicase and the RNA exosome to potentially exert control over RNA surveillance, and that this occurs in the vicinity of the ribosome. Proteomic landscapes and direct protein-protein interaction analyses demonstrated that aRNase J interplay with ASH-Ski2 and the Csl4 cap exosome subunit. These *in vitro* data are strengthened by our phylogenomic studies showing a taxonomic co-distribution of aRNase J and ASH-Ski2 among the archaeal phylogeny. Finally, our *T. barophilus* whole-cell extract fractionation experiments provide evidences that an aRNase J/ASH-Ski2 complex might exist *in vivo* and hint at an association of aRNase J with the ribosome or polysomes that is stressed in absence of ASH-Ski2. While aRNase J homologues are found among bacteria, the RNA exosome and the Ski2-like RNA helicase have eukaryotic homologues, underlining the mosaic aspect of archaeal RNA machines. Altogether, these results suggest, for the first time, a fundamental role of β-CASP RNase/helicase complex in archaeal RNA metabolism. Finally, our results position aRNase J at the junction of RNA surveillance and translation processes, thus opening new perspectives and evolutionary scenario on RNA processing players in Archaea.

## INTRODUCTION

Post-transcriptional regulation of gene expression demands accurate and timely RNA processing and decay to ensure coordinated cellular behaviours and fate decisions. Therefore, understanding RNA metabolic pathways and identifying RNA processing machineries, composed in general of ribonucleases (RNases) and ancillary enzymes such as RNA helicases, are major challenges in RNA biology. Currently, the best-understood RNA-dedicated pathways at the molecular level are those of Bacteria and Eukarya. In contrast, in Archaea, these molecular processes have been overlooked and are far from being understood.

Archaea, micro-organisms with signature sequences reported in all terrestrial and in human microbiome (1), have attracted considerable attention because of the orthologous relationships existing between their information processing machineries and those of eukaryotes (2–8). Regarding RNA machineries, it is worth mentioning that the archaeal 70S ribosome appears to be a simplified orthologous version of the eukaryal 80S ribosome with a reduced number of protein (9, 10) and that the archaeal RNA polymerase (RNAP) shares several features with eukaryal RNAPII such as similarities in amino acid sequences and of structures (11, 12). Furthermore, most archaeal genomes contain genes encoding an evolutionary-conserved phosphorolytic 3’-5’ RNA-degrading machinery, the RNA exosome (13), with the exception of Halophiles and some methanogens that possess homologues of bacterial RNase R (14, 15). In addition to its ribonucleolytic activity, the archaeal RNA exosome possesses a 3’-end RNA-tailing activity (14). This machinery, which shares high sequence and structure similarity with its eukaryotic counterpart, is constituted of a central catalytic core of three dimers of Rrp41-Rrp42 forming an hexamer ring with the three Rrp41 subunits carrying the catalytic activity (16–18). On the top, an RNA-binding platform composed of a trimer of Rrp4 and/or Csl4 subunits that has high affinity for A-rich RNA sequences and that is required for effective RNA degradation form the RNA exosome cap (19–22). In addition, a protein annotated as DnaG, composed of a primase domain, is part of the RNA exosome cap through an interaction with Csl4 (23). To date, the contribution of the archaeal RNA exosome to specific biological pathways is still unknown and it remains to determine if archaeal cells harbour specialized RNA exosomes, with heterogeneous and/or homogenous trimer cap composition *in vivo*.

The β-CASP RNases appear as versatile ancient enzymes with dual endo- and 5’-3’ exo-ribonucleolytic activities that act to control RNA maturation and stability in Bacteria and Eukarya (24, 25). Our previous studies identified and characterized three major groups of archaeal β-CASP RNases: aCPSF1 and aCPSF2 orthologous to the eukaryal Cleavage & Polyadenylation Specific factor CPSF73 and aRNase J orthologous to bacterial RNase J (26). This composite setting raises the question of the role of each β-CASP group in archaeal RNA homeostasis as well as the evolutionary origin of this family of enzyme among the three domains of life. In recent highlights on this mosaic system, we proposed potential functions within RNA-degradation machineries in Archaea (15, 27).

In Eukarya and Bacteria, as a general common thread, both mRNA decay and translation are intimately coordinated. These involve, after decapping/deprotection and deanylation, the 5’-3’ exoribonucleolytic activity supplied by the eukaryal Xrn1 or bacterial RNase J exo-RNases, which is complemented by the 3’-5’ exoribonucleolytic activity supplied by the eukaryal RNA exosome or bacterial PNPase, respectively (28, 29). Therefore, conserved 5’-3’ exo-ribonucleolytic activities are crucial in mRNA homeostasis in all eukaryotic and most bacterial cells. In archaeal cells, the 5’-3’ exoribonucleolytic activity is also conserved and is carried by aRNase J and aCPSF2. The role of this activity has been little studied (30, 31) and it remains to be clarified in which specific RNA decay and processing pathways it is involved. aRNase J and aCPSF2 exo-RNases have been identified being highly processive and having a preference for mono-phosphorylated RNA substrates in Euryarchaeota and Crenarchaeota, respectively (32, 33). In addition, a directional 5’-3’ RNA decay pathway was proposed to be at play in the crenarchaeal *S. solfataricus* cells. *In vitro* and *in vivo* studies evidenced cap-like structure of translation initiation factor (aIF2-γ), protecting RNA 5’-triphosphorylated ends from a 5’-end-dependent decay (30, 33, 34).

In this study, we focus on aRNase J as we identified this enzyme to be encoded in most of the Euryarchaeota phylum (26, 32, 35, 36). Towards understanding its physiological functions and relevance of 5’-3’ ribonucleolytic activity of the aRNase J is a mandatory step towards understanding its impact on cellular RNA homeostasis and in deciphering its exact tasks in RNA-maturation and decay pathways in euryarchaeal cells. Using combined biochemical, proteomics and genetics coupled with phylogenomic analyses, we present evidences that, in Thermococcales cells, the aRNase J cross-talks with the RNA exosome and we show that aRNase J forms a widely-conserved multi-protein complex with the archaeal specific helicase of the Ski2 family, ASH-Ski2. Moreover, our mutational analyses pinpoint functionally important domains involved in the aRNase J/ASH-Ski2 and aRNase J/Csl4 protein-protein interactions, respectively. Finally, we observed that a potential aRNase J-ribosome interaction exists and that could physically link 5’-3’ mRNA decay to translation in Euryarchaeota.

Altogether, our work builds the first blocks of complexes and networks involved in RNA-metabolic pathways in Archaea as we propose that aRNase J participates in RNA decay routes in the vicinity of the ribosome, in coordination with the ASH-Ski2 helicase and/or the RNA exosome, pointing a close relationship between the RNA exosome and Ski2-like helicase in Archaea. The mosaic setting of players around the 5’-3’ exo-RNase aRNase J, which is orthologous to bacterial RNase J, give a milestone towards the conservation of general principles of RNA-processing across the three domains of life since the ASH-Ski2 and RNA exosome have homologues found in Eukarya.

## RESULTS

### The protein-protein interaction network of aRNase J includes key components of RNA metabolism

To explore the protein interaction network of the euryarchaeal aRNase J, we carried out affinity purification coupled to mass spectrometry (AP-MS) analyses in *Pyrococcus abyssi* cell extracts, as already described (37). Recombinant aRNase J from *P. abyssi,* with C-terminal (His)_6_-tag (aRNase J-His), was produced, purified and used as baits for AP-MS experiments. In order to discriminate specific protein interaction from column background, we performed mock AP-MS analyses with *P. abyssi* cell extracts in absence of protein bait. In addition, to determine if some interactions are mediated through RNA or DNA, we implemented a nuclease treatment after incubating the cell extract with the bait. The co-purified partners were identified by bottom-up proteomic techniques coupled with mass spectrometry. The most significant partners identified in triplicate AP-MS are shown on the graph of **Figure 1a**. The extensive list of partners establishing significant interactions is given in Appendix **Table S1**. Remarkably, we found that aRNase J is at the centre of a protein interaction network that includes proteins with central functions in physiological processes. A majority of the protein partners of aRNase J-His are annotated as involved in RNA metabolism from transcription, RNA modification to RNA decay (**Figure 1a**). Indeed, significant interactions were detected with DnaG, a cap-associated subunit of the 3’-5’ RNA exosome decay machinery, with TmcA, the tRNA(Met) cytidine acetyl transferase (38), with the Fibrillarin and Nop5p, in charge of the 2’-O-methyltransferase activity of the C/D box guide RNP complex (39, 40) and with PAB2313, a putative ATP-dependent RNA helicase (**Figure 1a**). Remarkably, in addition to DnaG, the Rrp41 and Rrp42 catalytic core subunits and the Rrp4 cap subunit of the RNA exosome were also pull-downed. The subunits of the RNA polymerase (RpoA1/A2) and, one protein of the large ribosomal subunit particle were also observed. Thus, in light of these results, we propose that the 5’-3’ exonuclease aRNase J interacts with the most noteworthy RNA machineries of the cell.

**FIG. 1.**
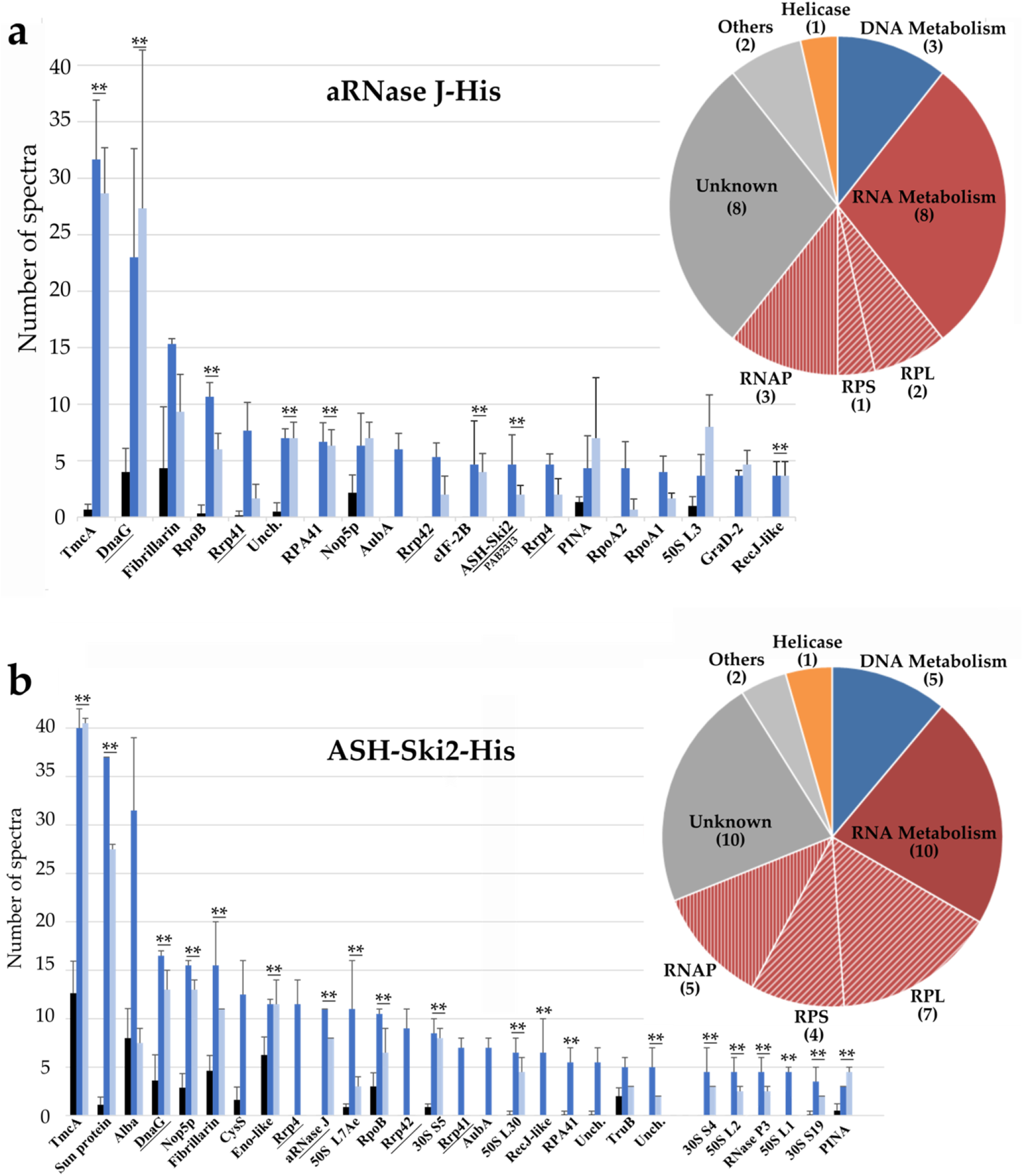
The most significant protein partners pull-downed with recombinant aRNase J-His in **a** and ASH-Ski2-His in **b** from whole-cell extract of *P. abyssi*. The histogram plots highlight the number of spectra obtained by mass spectrometry (Y-axis). The mean and standard deviation are shown. **Statistically significant difference verified by Student’s t-test (P < 0.01). Protein spectra from pull down assays carried out with or without nuclease treatment are indicated by light blue or darker blue bars, respectively. Protein spectra from the control assay in absence of the bait protein are signified by black bars. Only candidate with a number of spectra equal or higher than 5 in either condition was represented. For comprehensive lists of putative protein partners, see Appendix Tabl. S1 and S2A, respectively. The aRNase J, the RNA exosome subunits and the Ski2-like protein ASH-Ski2 (PAB2313) are underlined. Accordingly to the genome annotation of *P. abyssi*, the pie charts illustrate the proportion of protein partner with functions in RNA metabolism (red quotas), in DNA metabolism (blue quotas) or with putative helicase activity (yellow quotas).

In addition, proteins known to be involved in DNA maintenance processes, like RPA41, were also retrieved (**Figure 1a;**Appendix **Table S1**). RPA41 corresponds to the large subunit of the RPA complex that binds to single-stranded DNA. Interestingly, a physical interaction between RPA41 and the RNA polymerase has been previously demonstrated in *P. abyssi* (37). It was revealed that the RPA complex enhance transcription rate *in vitro*, probably *via* RPA41 interacting with the non-template strand of the elongating complex. In this light, the presence of RPA subunit in the interaction network of aRNase J could be considered as a further evidence for the involvement of aRNase J in RNA metabolism. Finally, it should be noted that the endo-RNase aCPSF1, the other β-CASP enzyme encoded in *P. abyssi* genome (26), was not pull-downed in those experiments meaning that aRNase J and aCPSF1 RNases do not appear to cross-talk.

Furthermore, we carried-out AP-MS analyses using as bait His-aRNase J, a recombinant aRNase J from *P. abyssi* with an N-terminal (His)_6_-tag. Out of the 22 partners pull-downed with aRNase J-His, only three proteins were significantly recovered with His-aRNase J. These proteins are PAB2313, RPA32 and RpoA2. We believe that the location of the tag at the N-terminus interferes with most protein-protein interactions.

### The *P. abyssi* ASH-Ski2 helicase shares a common network with *P. abyssi* aRNase J RNase and the RNA exosome

Interestingly, in the pull-down of aRNase J, we retrieved PAB2313, a putative RNA helicase, that appears to be a member of the Ski2-like helicase family (41) renowned for their crucial role in RNA decay in eukaryotic cells (42) and we named it ASH-Ski2. To gain further experimental evidence of a coordinated action of aRNase J and ASH-Ski2, we performed new rounds of AP-MS analyses with recombinant C- and N-terminal (His)_6_-tagged versions of *P. abyssi* ASH-Ski2 as baits and obtained an exhaustive list of partners (Appendix **Tables S2a & S2b)**. Remarkably, aRNase J was detected as an interacting partner of both tagged-versions of ASH-Ski2 (**Figure 1b**; Appendix **Figure S1)**. Furthermore, putative partners with a similar function distribution as for *Pab*-aRNase J were identified. This includes important RNA dedicated enzymes and machineries such as, the RNA modification enzyme (TmcA), the four components of the RNA exosome and the C/D guide RNP machinery in charge of rRNA and tRNA 2’O-methylations (**Figure 1b**; Appendix **Figure S1**). This clearly shows that ASH-Ski2 and aRNase J networks are interconnected as schematized in **Figure 2a**.

**FIG. 2.**
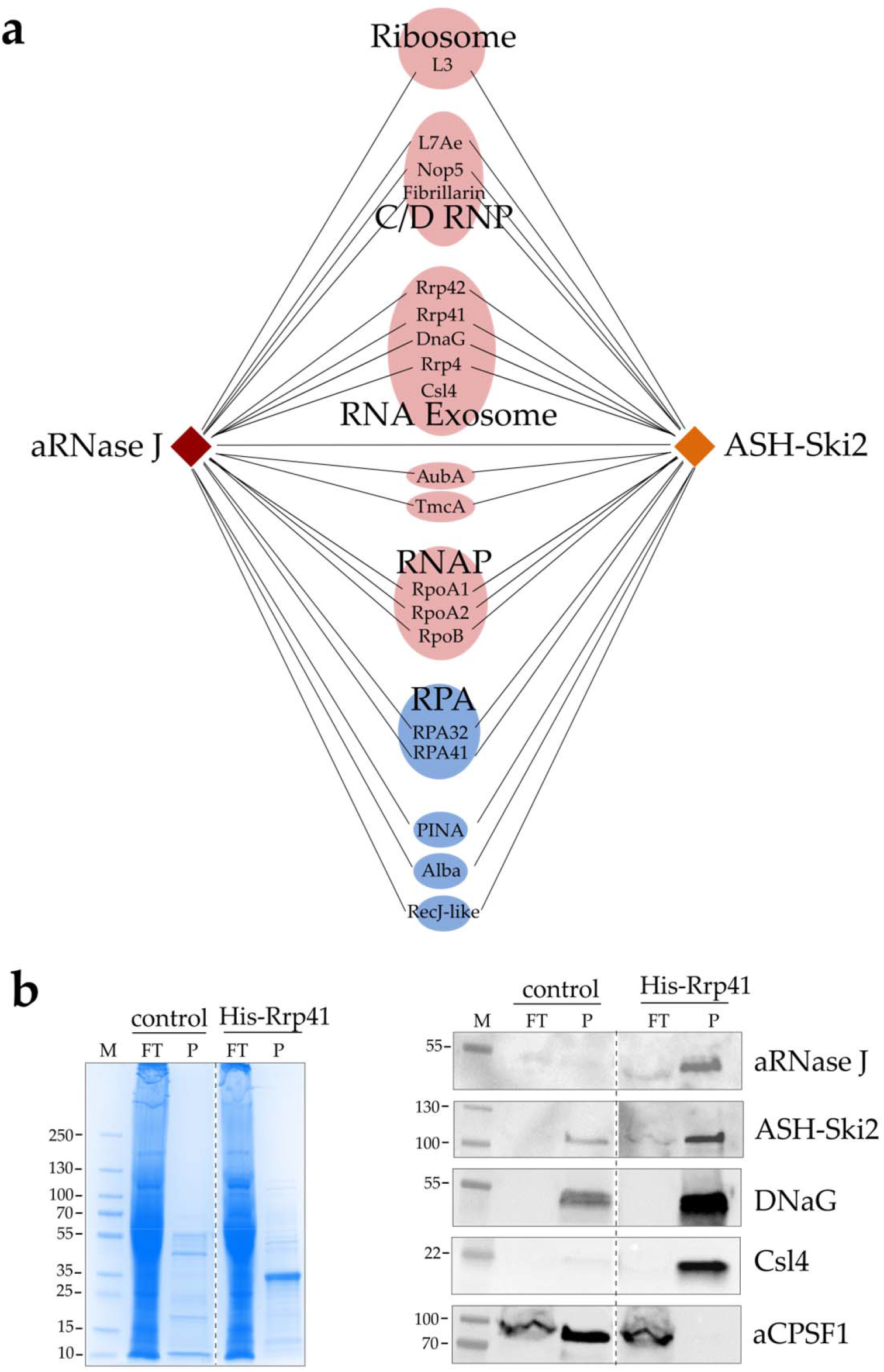
**a** Scheme representation of the protein partners shared by aRNase J-His (brown) and ASH-Ski2-His (yellow). **b** Pull down assays using the recombinant *Pab*-(His)-Rrp41 protein as bait in whole-cell extracts of *P. abyssi*; FT corresponds to the flow through, P to the pull-downed fraction and M to the protein ladder. Each fraction was analysed by coomassie blue staining of SDS-PAGE (left panel) and by Western blotting using specific antibodies for aRNase J, ASH-Ski2, DNaG, Csl4 and aCPSF1.

To ensure that aRNase J and ASH-Ski2 are part of the protein interaction network of the RNA exosome, we performed additional pull-down assays using a purified N-tagged His-Rrp41 of *P. abyssi* as bait. As before, a nuclease treatment was implemented to eliminate most of the interactions that would be mediated through RNA or DNA molecules. Among the pull-downed proteins, by western blotting, we specifically detected endogenous aRNase J, ASH-Ski2, Csl4 and DnaG but not the endo-RNase aCPSF1 (**Figure 2b**). Altogether, these results support a tight cross-talk between aRNase J, ASH-Ski2 and the components of the archaeal RNA exosome.

### The archaeal ASH-Ski2 members share identical taxonomic distribution as aRNase J members

To gain insights into the relationship that exists between archaeal aRNase J and ASH-Ski2 members as well as with the RNA exosome subunits, we compared their taxonomic distributions among the archaeal phylogeny. To do so, we first upgraded a phylogenomic study of the superfamily of the archaeal SF2 helicase (41) in order to accurately identify the Ski2 family of helicases encoded in archaeal genomes. In contrast to the role of eukaryotic Ski2-like RNA helicases in RNA homeostasis, reports on the functions of archaeal Ski2-like helicase family members are still scarce. As a common theme throughout SF2 helicase families, the unique characteristics of Ski2-like family members are mainly derived from accessory domains that decorate the helicase core and provide additional functionalities (43, 44). Ski2-like helicase structures demonstrate that the molecular “core” of all Ski2-like helicases is a ring-like four domain assembly of two RecA domains, a winged helix domain and a ratchet domain (42, 45). By performing RPS Blast searches, we retrieved all archaeal Ski2-like helicase member sequences from 147 non-redundant completely-sequenced archaeal genomes. In addition to Hel308 members, characterized by the COG 1204 and documented with enzymatic features of DNA helicases (42, 46–49), we also retrieved the ASH-Ski2 group, characterized by the COG 1202 (41). Interestingly ASH-Ski2 members hold an additional N-terminal domain in which four strictly conserved cysteines have the propensity to form a zinc-finger-like motif (50) (**Figure 3a**). ASH-Ski2 members are specific of euryarchaeal phylogenetic groups with the Lokiarchaeum as an exception (**Figure 3b**). Remarkably aRNase J and ASH-Ski2 members present similar occurrence, which is restricted to most euryarchaeal groups, with the exception of the archeoglobales and thermoplasmatales. Furthermore, the co-evolution of aRNase J and ASH-Ski2 members shown by the comprehensive congruence of aRNase J and ASH-Ski2 phylogenetic trees, with only few observed leaf displacements within taxonomic groups, reinforces their potential functional interplay (Appendix **Figure S2**). Altogether, protein networks combined with our phylogenomics analyses argue for a coordinated action of aRNase J and ASH-Ski2 in specific cellular processes in Euryarchaea.

**FIG. 3.**
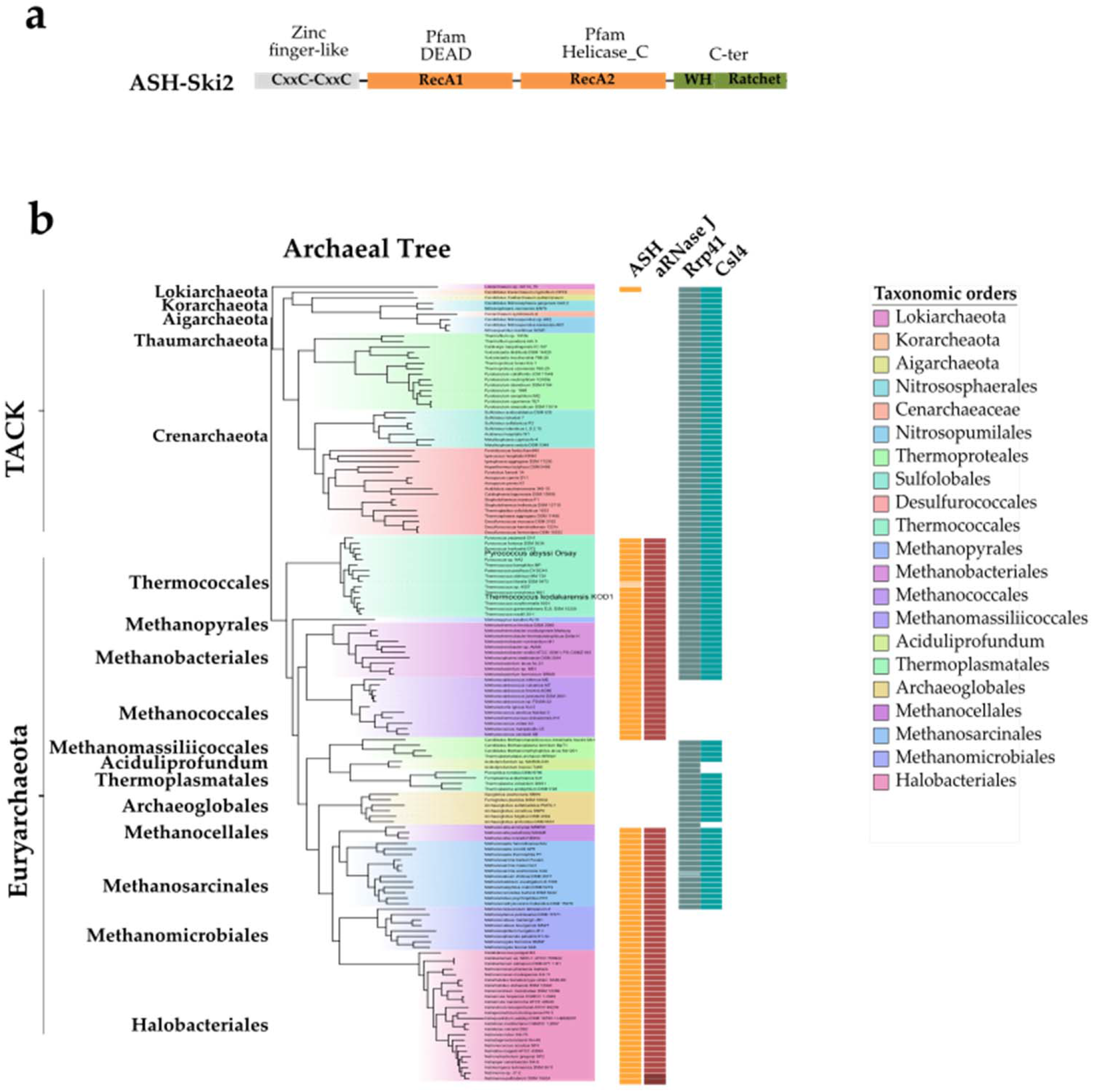
**a** Schematic protein domain architecture of the ASH-Ski2 group. The common SF2 core is composed of the RecA1 and RecA2 subdomains (Pfam DEAD & Helicase_C) in orange, of the C-terminal domain in green and of the N-terminal domain in grey. The strictly conserved cysteine residues of the N-terminal domain are represented. **b** Taxonomic distributions of ASH-Ski2, aRNase J and Rrp41/Csl4 from the RNA exosome among archaeal genomes. Their occurrence is plotted and juxtaposed to the archaeal specie tree. Taxonomic orders and Pfam domains are colour coded. Darker and lighter shades indicate multiple copies and pseudogenes, respectively.

Subsequently, we compared the taxonomic distribution of aRNase J, ASH-Ski2 and the RNA exosome Rrp41 and Csl4 subunits along the archaeal phylogeny (**Figure 3b**). Like in our previous studies (15, 27), we showed that the RNA exosome is not present throughout the whole archaeal taxonomy and that aRNase J, as well as ASH-Ski2, do not follow the taxonomic distribution of RNA exosome components. Indeed, it is absent from the Halobacteria and Methanomicrobiales members for which the 3’-5’ exo-ribonucleolytic activity is carried by another enzyme, the aRNase R (14, 15). In the Methanococci group, no enzyme with equivalent activity could be, so far, detected by comparative genomics.

### aRNase J forms a stable complex with the helicase-like ASH-Ski2, *in vitro*

To investigate a direct interaction between aRNase J and ASH-Ski2, we used an *in vitro* co-purification assay on nickel affinity chromatography to test pairwise interactions. A recombinant (His)_6_-tagged bait (His-ASH-Ski2) and an untagged prey (aRNase J) from *P. abyssi* were expressed independently in *E. coli* cells using the pET15b and pET11b expression vectors, respectively (Appendix **Table S3**). Prey and bait cellular extracts were mixed prior the lysis step. After nucleic acids removal with a cocktail of nucleases, the thermosensitive proteins from *E. coli* were eliminated at 70°C for a specific enrichment of the thermo-resistant proteins from *P. abyssi*. The clarified cellular extracts were then injected on a nickel column to retain the bait His-ASH-Ski2 protein and its potential interacting partners. Washing steps with low imidazole concentrations served at removing weakly bound contaminants. The bait and the retained interacting proteins were eluted with a linear gradient of imidazole. The protein content of each recovered fractions was analyzed by SDS-PAGE. The specific presence of aRNase J and His-ASH-Ski2 was revealed by Western Blotting using specific antibodies as shown in **Figure 4a**. As expected, aRNase J co-elutes with His-ASH-Ski2 even after nucleic acids removal. This establishes that aRNase J and His-ASH-Ski2 associate to form a stable complex that is not mediated through RNA or DNA molecules (**Figure 4a**,top panel). As a control, to demonstrate that protein co-purification is a consequence of complex formation rather than non-specific interactions with the column matrices, we controlled that untagged versions of aRNase J or ASH-Ski2 are not intrinsically retained on Ni^2+^-NTA column matrices (Appendix **Figure S3a**).

**FIG. 4.**
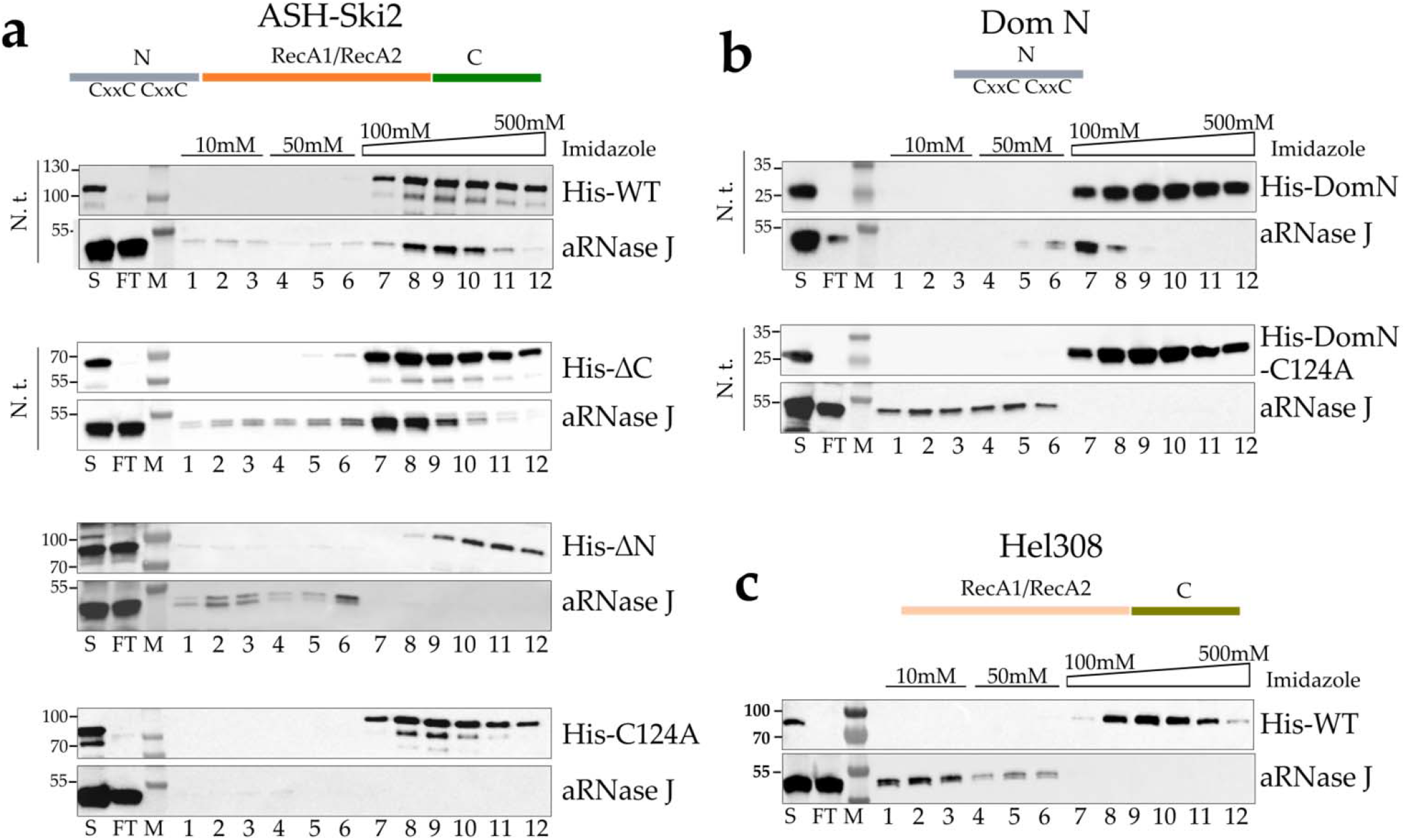
*In vitro* co-purification assay of aRNase J challenged by *Pab*-ASH-Ski2 and *Pab*-Hel308. **a** Schematic representation of the protein domain architecture of ASH-Ski2. Affinity co-purification assays on nickel chromatography column of *Pab*-(His)6-ASH-Ski2 and its variants. Recombinant full-length protein (WT), deleted for the C-terminal (ΔC) and N-terminal (ΔN) domains or harbouring a punctual substitution of the conserved cysteine residue into alanine (C124A) were tested. Proteins of each fraction were separated on a 4-15% SDS-polyacrylamide gel and the presence of proteins was monitored by Western blot using specific antibodies. S: Supernatant loaded on the affinity Nickel column, FT: flow through, Wash steps with 10mM (lanes 1 to 3) and 50mM (lanes 4 to 6) imidazole Elution with an Imidazole gradient from 100 to 500mM (lane 7 to 12). These experiments were performed with or without nucleic acid treatment (N.t.) as indicated on the left of each panel. **b** Identical assays as in A with the N-terminal domain (DomN) of *Pab-*ASH-Ski2. **c**Identical assays as in **a** with *Pab*-Hel308.

To uncover which protein domain of ASH-Ski2 is specifically involved in the formation of a complex with aRNase J, variants of His-ASH-Ski2 were used as baits for the untagged prey aRNase J. Pairwise interactions were tested as previously described performing *in vitro* co-purification assays (**Figure 4a**,lower panels). Our results indicate that the accessory C-terminal domain of ASH-Ski2 (His-ΔC) is not required in establishing an *in vitro* interaction with aRNase J as the elution profile is comparable to the one with full-length ASH-Ski2. In contrast, the accessory N-terminal domain of ASH-Ski2 (His-ΔN) seems to be critical for the interaction since aRNase J does not co-elute with the ΔN variant of ASH-Ski2. Finally, we showed that the accessory N-terminal domain of ASH-Ski2 expressed alone (His-DomN) is sufficient to establish an *in vitro* interaction with aRNase J (**Figure 4b**,top panel).

The sequence alignment of the N-terminal domains of archaeal ASH-Ski2 highlights the presence of four strictly conserved cysteines with the propensity to form a zinc finger-like motif (**Figure 3a**). To test whether this motif is critical for the interaction between ASH-Ski2 and aRNase J, we substituted the C124 residue of full-length His-ASH-Ski2 and His-DomN by an alanine (A), and we challenged their capacity to retain aRNase J on on Ni^2+^-NTA column as before. Our results show that the conserved C124 residue is critical in establishing or in stabilizing an interaction between the full-length or DomN of ASH-Ski2 and aRNase J (**Figure 4a & 4b, lower panels)**. Indeed, the single C124A substitution is sufficient in destabilizing this interaction

Finally, to confirm the specificity of the interaction between aRNase J and ASH-Ski2, we performed similar *in vitro* co-purification assays using aRNase J as a prey and His-Hel308, another Ski2-like helicase encoded in the *P. abyssi* genome, as bait. In this configuration, aRNase J is not retained on the Ni^2+^-NTA column matrices (**Figure 4c**) meaning that aRNase J specifically interacts with ASH-Ski2. This result is not surprising as members of the Hel308 family do not possess the N-terminal zinc finger-like motif extension.

### aRNase J forms a stable complex with the exosome cap-subunit Csl4 *in vitro*

We also tested the pairwise interactions between aRNase J or ASH-Ski2 and RNA exosome subunits that are among the most retrieved proteins with aRNase J and ASH-Ski2 in the AP-MS analyses (**Figure 1 & Figure 2**). The results of the *in vitro* co-purification assays performed as described above are summarized in **Figure 5a**. First, we challenged the catalytic subunit His-Rrp41 as bait with aRNase J or ASH-Ski2 as preys. Only aRNase J could be recovered in the elution fractions together with His-Rrp41. The Rrp41/aRNase J complex is nucleic acid-dependent since its formation is sensitive to nuclease treatment (Appendix **Figure S3b**). Then, we challenged the exosome cap subunits His-Rrp4, DnaG-His or His-Csl4 as baits with aRNase J or ASH-Ski2 as preys. In our conditions, neither Rrp4 nor DnaG were found to interact with aRNase J or ASH-Ski2 (Appendix **Figure S3c**). However, we could detect an interaction between ASH-Ski2 and Csl4, but this association appears to be labile and sensitive to nuclease treatment (**Figure 5b**). In contrast, the exosome cap-subunit Csl4 is able to form a stable complex with aRNase J that is not sensitive to nuclease treatment and therefore is not mediated by nucleic acids (**Figure 5c**). To go further in the characterization of this association, we then searched for the protein domains of Csl4 that are involved in the formation of the Csl4/aRNase J complex. While the N-terminal domain of Csl4 (ΔN) is not required, its C-terminal domain (ΔC) is critical for the formation of the Csl4/aRNase J complex (**Figure 5c**).

**FIG. 5.**
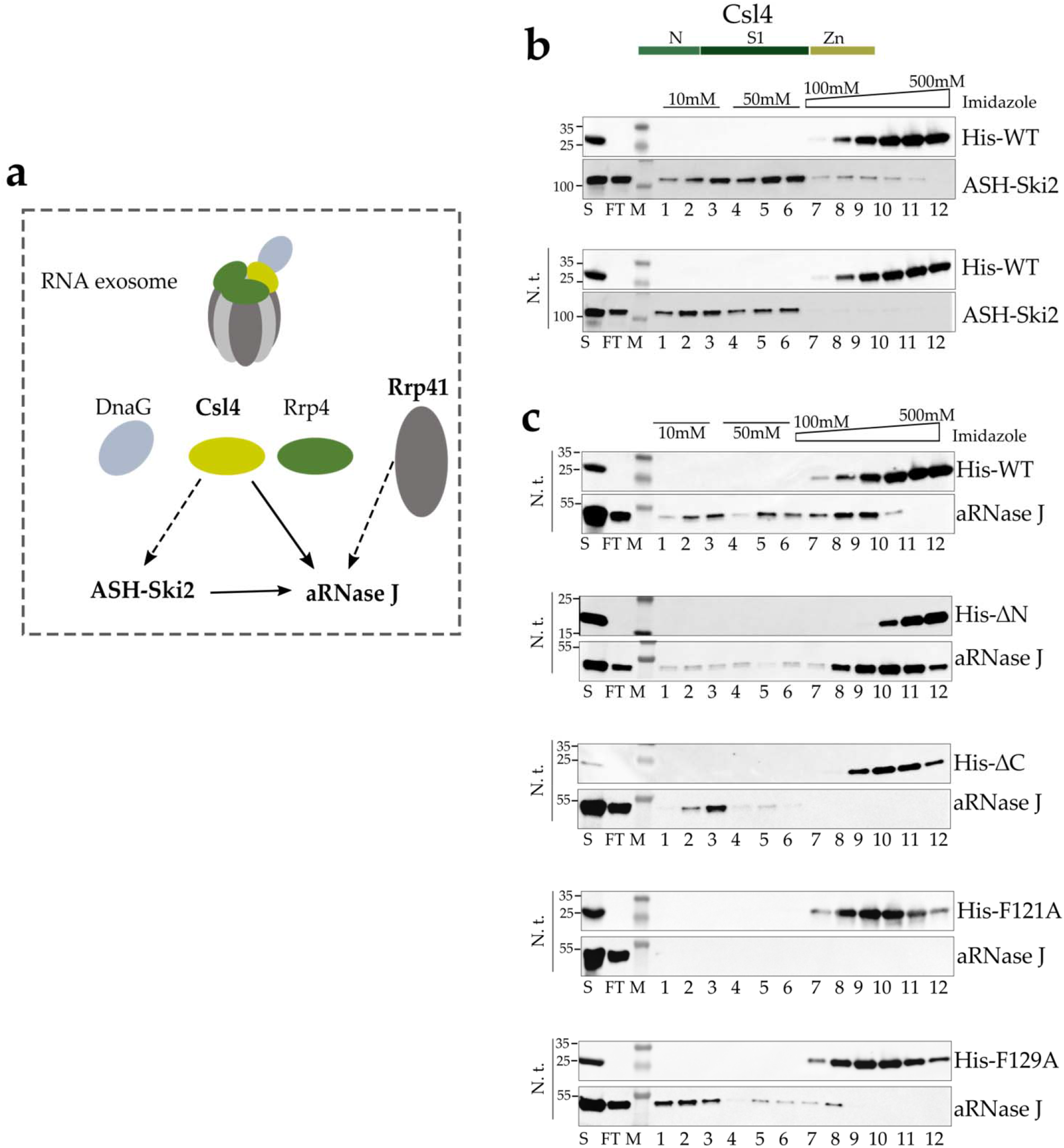
*In vitro* co-purification of *Pab*-aRNase J challenged by *P. abyssi* RNA exosome subunits. **a** Schematic representation of the RNA exosome machine and of *in vitro* protein interactions detected in this study. Plain arrows indicated an interaction resistant to a nuclease treatment. Dotted arrows indicate interaction sensitive to a nuclease treatment. he domain architecture of cap or core archaeal exosome subunits are shown (left). **b***Pab*-(His)-Csl4 was challenged *Pab*-ASH-Ski2. **c** The schematic representations of the protein domain architecture of *Pab*-(His)-Csl4 is shown *Pab*-(His)-Csl4 and its variants were challenged with *Pab*-aRNase J. Variants with a deletion of the C-terminal (ΔC) or N-terminal (ΔN) or a punctual substitution at the phenylalanine 121 and 129 residues (F121A, F129A) in alanine was tested. Legend as in Fig. 4.

To highlight specifically conserved residues in Csl4 sequences that are co-distributed with aRNase J, we derived Weblogos from multiple alignments of Csl4 co-distributed or not with aRNase J. Several positions (including F/Y121 and F129, numbering according to the *P. abyssi* sequence) in the S1 domain of Csl4 could be identified as highly conserved in Csl4 sequences in the context of the co-distribution with aRNase J (Appendix **Figure S4a**). Strikingly, the phenylalanine residues, F121 and F129, located on the central S1 domain of Csl4 and solvent-exposed in the *P. abyssi* Csl4 structural model (Appendix **Figure S4b)**are also key for the association with aRNase J (**Figure 5c**).

### aRNase J and ASH-Ski2 and the RNA exosome co-sediment in sucrose gradient fractionation of whole-cell extract from *P. abyssi and Thermococcus barophilus*

Altogether, our results propose a tight relationship between aRNase J, ASH-Ski2 and the RNA exosome. Here, we asked whether these protein partners could form higher order protein complexes in archaeal cells. To this end, we discriminated the sedimentation profile behaviours of endogenous aRNase J, ASH-Ski2 and Rrp41 in two Thermococcales organisms, *Thermococcus barophilus* (*Tba*) and *P. abyssi* (*Pab*) by doing ultracentrifugation of whole-cell extracts on continuous 10-30% sucrose density gradient (**Figure 6a**; Appendix **Figure S5a**, respectively). Since to this date, *P. abyssi* cannot be genetically modified, the *T. barophilus* strain was used as a genetically tractable model (51, 52) to assess the sedimentation profile of endogenous aRNase J in absence of ASH-Ski2 and vice-versa. Moreover, note that the antibodies produced to detect our proteins of interest from *P. abyssi* by immunoblotting cross-react with their counterparts from *T. barophilus* while preserving their specificity. Surprisingly, the vast majority of these proteins co-sediment with ribosomal fractions with barely any in light fractions. This is not the case for the β-CASP endo-RNase *Pab*-aCPSF1, which was not identified in any of the protein interaction networks that we uncovered in this study (Appendix **Figure S5).**It is interesting to note that *Pab*/*Tba*-aRNase J and *Pab*/*Tba*-ASH-Ski2 are similarly distributed through the gradient (**Figure 6a;**Appendix **Figure S5**). This distribution pattern is also highly similar to the one of the RNA exosome subunit *Pab*/*Tba*-Rrp41 (**Figure 6a &** Appendix **Figure S5**). In conclusion, our data are indicative of a co-migration of the *Pab*/*Tba*-aRNase J, the *Pab*/*Tba*-ASH-Ski2 and the Rrp41 RNA exosome subunit which follow ribosomal RNA species.

**FIG. 6.**
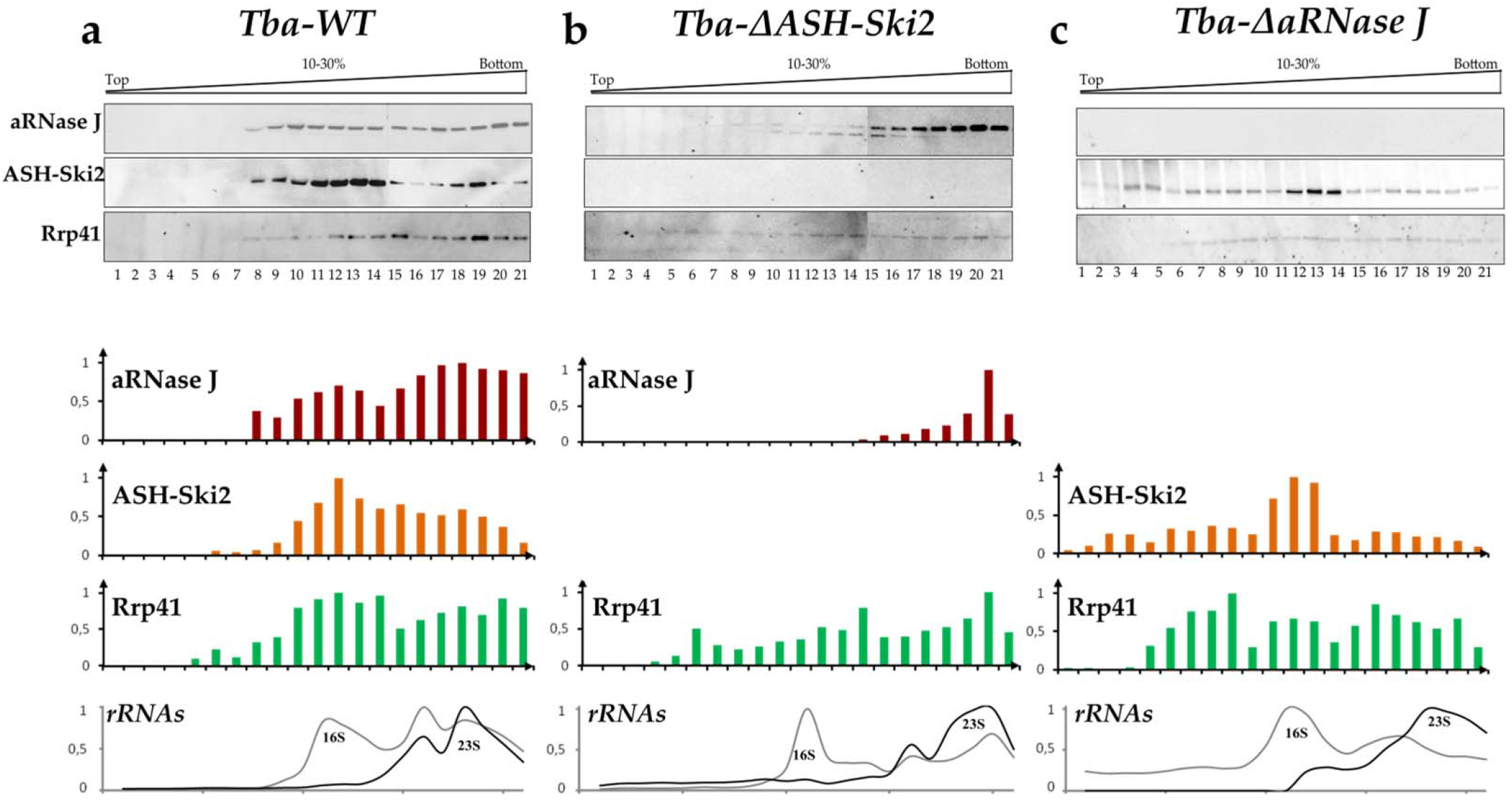
Sedimentation profiles of endogenous aRNase J, ASH-Ski2, Rrp41 and DnaG of wild type (WT) in **a**, ΔASH-Ski2 in **b**, and ΔaRNase J in **c**, *T. barophilus* strains. Clarified whole-cell extracts were fractionated on 10-30% density sucrose gradients by ultracentrifugation. Equal volumes of fraction, precipitated with TCA, were monitored by Western blotting using specific antibodies. The digital images and bar charts representing the relative amounts of protein are given. The ribosomal RNAs were also profiled by slot blotting equal volume of denatured fractions with probes complementary to 16S and 23S rRNAs. The relative quantification of 16S and 23S ribosomal RNAs is plotted in grey and black, respectively.

To go further, we fractionated whole-cell extracts from *T. barophilus* strains enclosing a deletion of the gene encoding *Tba*-ASH-Ski2 (*Tba-ΔASH-Ski2,* **Figures 6b**) or *Tba-*aRNase J (*Tba-ΔaRNase J*, **Figures 6C**). On one hand, when *Tba*-ASH-Ski2 is missing, most of the endogenous *Tba*-aRNase J protein is shifted towards the polysome fractions at the bottom of the gradient. On the other hand, when *Tba*-aRNase J is not present, the distribution profile of the endogenous *Tba*-ASH-Ski2 is also clearly altered; but in this case, *Tba*-ASH-Ski2 is more evenly spread out along the gradient with a main peak overlapping the 16S ribosomal RNA profile. Note that the profile of the core RNA exosome subunit, Rrp41, was barely affected in both genomics contexts. Overall, these results strongly suggest that the aRNase J/ASH-Ski2 complex that was identified *in vitro* might exists *in vivo* in *T. barophilus* cells.

We also challenged the physical association of *Tba*-aRNase J and *Tba*-ASH-Ski2 with the ribosome subunits, by performing similar fractionation assays but at low magnesium concentration (TK-EDTA buffer) to dissociate the assembled ribosomes into individual subunits (53). Under these conditions, in the wild-type strain, endogenous *Tba*-aRNase J and *Tba*-ASH-Ski2 co-sediment with the 16S and 23S rRNAs from the small and large subunits, respectively (Appendix **Figure S6a**). Again, we have a perfect overlay of the sedimentation profiles of *Tba*-aRNase J and *Tba*-ASH-Ski2 as previously seen at a higher concentration of magnesium (**Figure 6a**; Appendix **Figure S6a**). Same results are observed in the context of *P. abyssi* (Appendix **Figure S5b**). Conversely to what we observed at high magnesium concentration (TK buffer, **Figures 6b & 6c**), the sedimentation profiles of *Tba*-Rrp41 seems to be slightly affected by the absence of *Tba*-ASH-Ski2 (Appendix **Figure S6b**) and *Tba*-aRNase J (Appendix **Figure S6c**). Nonetheless, it is interesting to note an obvious shift of *Tba*-RNase J from the bottom of the gradient (when wild type *T. barophilus* cell extract at high magnesium concentration are considered) into two peaks overlapping the profiles of the 16S and 23S rRNAs, respectively, in the context of the *Tba-ΔASH-Ski2* strain, at low magnesium condition (Appendix **Figure S6b**). From these results, we argue for an association of aRNase J with ribosome or polysome that is stressed in absence of ASH-Ski2.

## DISCUSSION

Elucidating the RNA processing machineries is of a critical importance to understand the molecular basis of RNA decay and processing in Euryarchaea. In this study, to our knowledge, we provide the first experimental evidences hinting at the existence of high-order β-CASP protein complex(es) in Thermococcales that would carry 5’-3’ & 3’-5’ exoribonucleolytic and helicase-like activities and that could play pivotal roles in archaeal RNA surveillance. First, we identified the interaction landscape of aRNase J from *P. abyssi*, a 5’ to 3’ β-CASP exoRNase (32). Candidate partners include, notably, the archaeal RNA exosome that exhibits 3’ to 5’ exoribonucleolytic and 3’ end-tailing activities (17), the ribosome and a Ski2-like helicase ASH-Ski2. Of high significance, we also found these proteins to be candidate partners in the interaction landscape of ASH-Ski2 and in the pull-down of the core exosome subunit Rrp41. In addition, we showed that aRNase J is able to interact, *in vitro,* with ASH-Ski2 and the exosome cap subunit Csl4. Our *in vitro* data are strengthened by our phylogenomic studies showing that, with only one exception, ASH-Ski2 is only encoded in archaeal genomes that carry a gene for aRNase J and that amino acids of Csl4, involved in the *in vitro* interaction with aRNase J, are only conserved in archaeal genomes where their encoding genes are co-distributed. Finally, our *T. barophilus* whole-cell extract fractionation experiments provide evidences that an aRNase J/ASH-Ski2 complex might exist *in vivo* as their sequential deletions modify the sedimentation profile of their protein partner. Our fractionation assays also hint at an association of aRNase J with the ribosome or polysomes that is stressed in absence of ASH-Ski2.

In conjunction with our previous demonstration that β-CASP RNases, ubiquitous in Archaea, must be key players in archaeal RNA metabolism (26, 27), our current data support a model whereby the β-CASP RNase, aRNase J, the ASH-Ski2-like helicase and the RNA exosome interplay in most euryarchaeal cells (**Figure 7**). It remains to determine how this interplay could impact the archaeal RNA metabolism. Since our data imply that ASH-Ski2 has an impact on aRNase J and vice versa, we favour a scenario where, at least, ASH-Ski2 and aRNase J have a functional connection. This is all the more so given that aRNase J and ASH-Ski2 show a taxonomic co-distribution among the archaeal phylogeny and nearly perfect phylogenetic tree congruence suggesting a co-evolution and potentially an ancient origin. Based on what is known about the role of these factors in Eukarya and Bacteria, we discuss what we believe to be the most probable scenarios. Evidently, further work will be needed to determine how these proteins alone or in complex shape the euryarchaeal RNA metabolism.

**FIG. 7.**
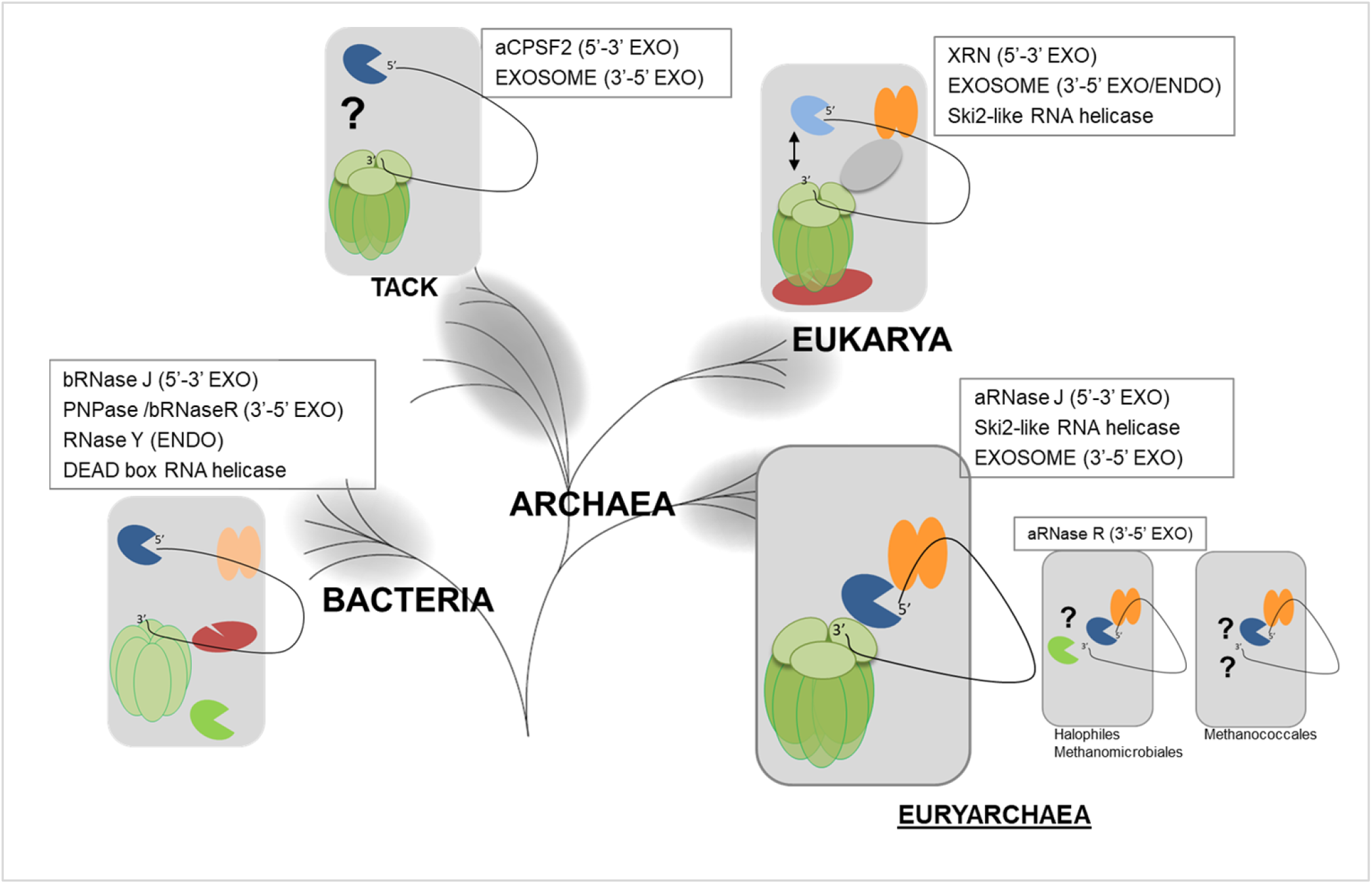
Schematic representation of the interplay between enzymes and machineries in charge of 5’-3’ and 3’-5’ RNA decay and surveillance in the three domains of life. The life tree model is as in (79). The 3’ end RNA trimming activity is carried by the RNA exosome machinery (barrel in green and cap in light green) in most Archaea (15) and Eukarya (61) that is similar to bacterial PNPase (barrel in light green). In Halobacteria and Methanomicrobiales the aRNase R, orthologous to bRNase R, (green packmans) carries the 3’-5’ exo activity. In Methanoccocci, no 3’-5’ exo activity has yet been described (15). The 5’-3’ exoribonucleolytic activities (blue packmans) are carried by XRN ribonuclease in Eurkarya (80), bRNase J and aRNase J in Bacteria and Euryarchaea and aCPSF2 in the TACK group. RNA helicase activities shown to be involved in RNA surveillance belong to the Ski2-Like family in Eukarya (42) and Euryarchaea (this study) and to the DEAD box family in Bacteria (81). The interplay between 5’-3’ and 3’-5’ exoribonucleolytic activities are reported to take place in the RNA-degradosome-like complex in Bacteria that also involves endoribonucleases (red packman) (82) and through the “closed-loop” architecture or 5’-3’ communication of mRNA in Eukarya (83). In most Euryarchaea,, we propose that aRNase J/ASH-Ski2 complex interplays with the RNA exosome. In the TACK group and the Halophiles, Methanomicrobiales and Methanoccocales, such an interplay remains to be characterized (question marks).

First, recent published work proposes that ASH-Ski2, also named Eta for euryarchaeal termination activity, influences transcription termination by disrupting archaeal transcription elongation complex (54). However, as its helicase activity is rather slow and cannot keep up with wild-type RNAP elongation rates, Eta was recently proposed to be more likely involved in the rescue of transcriptional arrest (55). In our study, the presence of RPA and RpoB subunits in the interaction network of ASH-Ski2 could be a further evidence for the involvement of ASH-Ski2 in the transcription. In addition, transcriptional termination and rescue pathways also involve proteins with ribonucleolytic activities. In Eukarya, the 5’-3’ exoribonucleolytic activity of Rat1p/Xrn2p was proposed to be associated in a transcription termination pathway called “torpedo model” (56). In chloroplast, RNase J was shown to play a critical role in removing antisense RNAs that are generated by read-through transcription (57).

Nevertheless, chloroplast RNase J exhibits a DNA binding domain found in transcription factors of plants that is not present in archaeal RNase J and that could be essential to fulfil this function (58). Finally, co-factors of the nuclear RNA exosome were shown to connect transcription termination to RNA processing by guiding terminated transcripts to the appropriate exonuclease (59). In Archaea, it is still not known if a ribonucleolytic activity is critical since, until now, the complete picture of transcription termination is far from being understood (55). It remains to identify if aRNase J, ASH-Ski2 and the RNA exosome are involved together or alone in transcription termination or rescue pathways which have been shown to be key in influencing the fate of a transcribed RNA.

Moreover, based on the roles of the Ski2-like helicases in Eukarya (60), the idea that the euryarchaeal ASH-Ski2 group is closely connected to RNA-related pathways emerges (**Figure 7**). Indeed, eukaryal Ski2, Mtr4 and Brr2 which are described as RNA helicases that process RNAs in a 3’-5’ direction were shown to play critical roles in RNA decay, surveillance and splicing, respectively (42, 60). Notably, Ski2 and Mtr4 are essential co-factors of the cytoplasmic and nuclear RNA exosomes, respectively, by being involved in the recruitment of RNA substrates (13). Likewise, our work provides several evidences of interplay between the archaeal RNA exosome and ASH-Ski2. Our data also suggest that this interplay is done through aRNase J that interacts *in vitro* with Csl4 from the RNA exosome cap and with ASH-Ski2. Altogether, our results converge towards a cross-talk between the 5’-3’ and 3’-5’ exo-ribonucleolytic activities carried by aRNase J and the RNA exosome, respectively. This is consistent with what is observed in Eukarya, where mounting evidences show a functional interplay between surveillance pathways taking place at the 5’ and 3’-ends of RNAs (61) (**Figure 7**).

Finally, it must be remembered that aRNase J is the homolog of bacterial RNase J (bRNase J) which was shown to be critical in mRNA decay and rRNA maturation pathways and to be part of a degradosome-like complex (29, 62, 63). bRNase J which can carry 5’-3’ exo- and endo-RNase activities was shown to associate with proteins identified as, among others, helicase-like proteins and a polynucleotide phosphorylase (PNPase). Bacterial PNPase form an RNA-degradation machine with 3’-5’exorinucleolytic and polyadenylation activities reminiscent to the archaeal RNA exosome (64) (**Figure 7**). Although, in some organisms, the bRNase J degradosome-like complex appears to be transient and dynamic, limiting its validation *in vitro*, this is consistent with interplay between 5’-3’ and 3’-5’ RNA surveillance decay in Bacteria. Therefore, it emerges that, in the three domains of life, RNA surveillance could be taken care of by evolutionary conserved RNase-based RNA-degradation machines with RNA helicase and with 5’-3’ and 3’-5’ exoribonucleolytic activities (**Figure 7**).

From cellular fractionation assays, we observed that both aRNase J and ASH-Ski2 could potentially associate with the ribosome. Interestingly, it was also shown that bRNase J can associate with translating ribosomes (65) and the crystal structure of the eukaryotic 80S ribosome-Xrn1 complex showed that the 5’-3’ Xrn1 nuclease binds at the mRNA exit site of the ribosome (66). Since the sedimentation profiles of *T. barophilus* and *P. abyssi* aRNase J resemble to the one reported for the eukaryal 5’-3’ exo-RNase Xrn1, we speculate that the archaeal aRNase J is also associated with polysomes in Thermococcales cells. This is reminiscent of what is described in eukaryal mRNA surveillance pathways (28) in which mRNA decay is initiated by an endonucleolytic cleavage of ribosome-associated mRNAs (67, 68), followed by rapid degradation in the 3′-5′ direction by the RNA exosome or in the 5′-3′ direction by Xrn1 (69). Recently, direct experimental evidence showed that the cytoplasmic RNA exosome is able to interact with the ribosome through a direct interaction mediated by the SKI complex (composed of Ski2-Ski3-Ski8 factors). More precisely, it was shown that ribosome binding displaces the auto-inhibitory domain of the Ski2 RNA helicase (70).

On a side note, it is important to notice that the RNA exosome is not present throughout the whole archaeal taxonomy (15, 27) and that aRNase J does not accurately follow the taxonomic distribution of RNA exosome components. In the Halophiles that do not encode the exosome subunits, the 3’-5’ exo-ribonucleolytic activity is carried by another enzyme, the aRNase R (14, 15). It would be interesting to know whether the halophilic and methanomicrobiale aRNase J, alone or in complex with ASH-Ski2 complex, interplays with aRNase R. In the Methanococci group, no 3’-5’ exo-RNase could be detected by comparative genomics. In this case, aRNase J might connect to specific archaeal RNases that remain to be identified (**Figure 7**). In conclusion, this work establishes a solid base in the largely overlooked but extremely important field of archaeal RNA metabolism. We identified two phylogenetical conserved factors in Euryarchaea, the 5’-3’ exo-RNase aRNase J and the Ski2-like helicase ASH-Ski2, which have homologs in Bacteria and Eukarya, respectively (**Figure 7**). Therefore, a mosaic system in Euryarchaea could be at the centre of networks interplaying with the archaeal RNA exosome and the ribosome. This is an example showing that Archaea use composite systems and thereby elucidating mechanism in archaeal organisms could give valuable insights into understanding function and evolutionary routes of protein networks or/and high-order complexes. Analogous to eukaryal and bacterial systems, we propose that these factors act in partnership in major archaeal RNA decay or surveillance pathways. Deeper insights need to be gained into the function and importance of these enzymes in mRNA decay, ribosomal RNA maturation, mRNA 3’-end formation and transcription termination. It remains, among other aspects, to be established how these factors might be involved in shaping the RNA landscape in euryarchaeal cells. Finally, the settlement of specific enzymatic properties of ASH-Ski2 and the determination of structure-function of ASH-Ski2/aRNase J and Csl4/aRNase J complexes will ascertain their physiological functions in the context of the interplay with aRNase J.

## MATERIAL & METHODS

### Vectors and Oligonucleotides

The supplementary Tables S3 and S4 summarize T7-promotor-driven pET vectors and oligonucleotides used in this study. All constructions were obtained by assembling PCR fragments using InFusion® cloning kit (Takara). Using appropriate sets of oligonucleotides, pET vectors were amplified with the PrimeSTAR Max DNA polymerase (Takara) and the coding sequence of *Pyrococcus abyssi* aRNase J (PAB1751), ASH-Ski2 (PAB2313), Hel308 (PAB0592), DnaG (PAB0316), Rrp41 (PAB0420), Rrp4 (PAB0419) and Csl4 (PAB2314) were amplified from genomic DNA using the Phusion High-Fidelity DNA polymerase (ThermoFisherScientific). The pET vectors expressing the ASH-C124A, Csl4-F121A and Csl4-F129A variants were generated by site-directed mutagenesis of their wildtype counterparts with appropriate sets of oligonucleotides using the QuikChange II XL Kit (Stratagene) as recommended.

### *T. barophilus* strains

The chromosomal copy of the TERMP_01768 (encoding *Tba*-ASH-Ski2) and the TERMP_00146 (encoding *Tba*-aRNase J) were deleted by the pop-in/pop-out method to generate the *Thermococcus barophilus* ΔASH-Ski2 and ΔaRNase J strains, respectively using published protocol (51, 52).

### Production and purification of bait proteins

*E. coli* BL21-CodonPlus (DE3) cells freshly transformed with pET15-aRNase J, pET15-ASH-Ski2, pET21-ASH-Ski2, pET15-Csl4 or pET15-Rrp41 vectors (Table S3) were grown in 400mL of LB medium at 37°C. Protein production was induced in exponential phase at an OD_600nm_ of 0.6 by the addition of 0.1mM IPTG. After 3h of induction at 30°C, the cells were collected, suspended in 10 mL of lysis buffer (50mM NaPhosphate, 300mM NaCl, 10mM Imidazole) supplemented with 1mg.mL^−1^ of lysozyme and a mix of EDTA-free protease inhibitor (cOmpleteTM, Roche) and lysed by sonication (4x[5*10s], 50% cycle, VibraCell Biolock Scientific). When mentioned, the cleared extracts, obtained by centrifuging the crude extracts (20,000g 4°C, 20min), were treated with a mix of RNase A (20μg.mL^−1^), RNase T1 (1U.μL^−1^) and DNase I (20μg.mL^−1^) containing 10mM of MgCl_2_ for 30min at 37°C. After a heating step at 70°C for 20min, the extracts were furthered clarified by centrifugation (20,000g, 4°C, 20min). The recombinant proteins were two-step purified from the soluble fractions to near homogeneity using FPLC (Fast Protein Liquid Chromatography, Äkta-purifier10, GE-Healthcare): first, by a nickel-attached HiTrap chelating column (HisTrap 1mL, GE-Healthcare) with a linear gradient of 50-500mM imidazole; secondly, by a heparin column (HiTrap Heparin 1ml, GE-Healthcare) with a linear gradient of 300mM to 1M NaCl. The eluted fractions containing His-tagged proteins were pooled and dialyzed overnight against 500mL of dialysis buffer (20mM HEPES pH 7.5, 300mM NaCl, 1mM DTT, 10% glycerol buffer).

### Co-purification by chromatography affinity

Cells expressing His-tagged bait proteins were produced as described above. Cells expressing untagged prey proteins were obtained after transforming *E. coli* BL21-CodonPlus (DE3) cells with pET11-aRNase J, pET11-ASH-Ski2, pET11-Csl4 or pET11-*Pab*-Rrp41 vectors (Table S3) and using the same conditions for growth and induction. Two cell pellets from 200mL of cultures expressing either bait or prey protein were mixed upon lysis (as before). The bait proteins alone or in complex with the prey proteins were purified by nickel affinity chromatography using Ni^2+^-NTA column matrices (HisTrap 1mL, GE-Healthcare). The elution was obtained with a linear 50mM-500mM imidazole gradient after washing steps at 10mM and 50mM of imidazole. For each couple of bait/prey that were tested, the co-purification assays were at least performed twice either in presence or absence of RNase/DNase treatment (as above). The proteins from recovered fractions were analysed by Western blotting (4-15% Mini PROTEAN® TGX Stain-Free™ Gels & Trans-blot® Turbo™ Nitrocellulose Transfer Pack, BioRad). The bait proteins were probed using a His Tag HRP-conjugated antibody (Ozyme) diluted 5,000-fold. The prey proteins were probed using polyclonal antibodies against *Pab*-aRNase J, *Pab*-ASH-Ski2, *Pab*-Csl4, *Pab*-Rrp41, *Pab*-DnaG or *Pab*-aCPSF1 (custom polyclonal antibodies, Eurogentec) diluted 10,000-fold and an anti-rabbit IgG HRP conjugate (Promega) diluted 5,000-fold.

### Pull-down by affinity purification assays

The pull-down assays were performed in triplicate with clarified whole-cells extract of *P. abyssi* cultivated on exponential growth phase in bioreactors under physiological conditions (95°C, pH 6.5, anaerobic). The cell extracts were prepared as described in (37). Briefly, *P. abyssi* pelleted cells were re-suspended in 1/3 w/v of PBS buffer (recipe), supplemented with 300mM of NaCl and a mix of EDTA-free protease inhibitor (cOmpleteTM, Roche). After sonication (VibraCell Biolock Scientific), the crude extracts were clarified by centrifugation (20,000g, 30min, 4°C).

Purified His-tagged proteins were used as baits in clarified whole-cell extract of *P. abyssi*. Briefly, 20μg of bait proteins were immobilized on 0.6mg of cobalt-coated magnetic beads (Dynabeads, Invitrogen). The complex baits-beads were further incubated with 2mg of *P. abyssi* extract under rotation for 2 hours at room temperature. The protein complexes formed *in vitro* were separated on a magnet and washed extensively with PBS buffer (3*4mL and 2*1mL). A second round of pull-down assays was performed with an additional step to eliminate non-specific interaction via DNA or RNA molecules by incubating the protein complexes with an RNase /DNase mix (10μg.mL^−1^ of RNase A and DNase I) for 30 min at room temperature before applying the magnetic force. A control was also performed under identical conditions using cobalt-coated beads alone instead of the baits-beads complexes. Purified protein complexes were eluted in 25μL of XT sample buffer (BioRad) containing 2μL of 20X reducing buffer at 95°C for 10 min and separated by SDS-PAGE short migration on 12% SDS-PAGE (Criterion XT Precast gels–BioRad). After a short migration, the proteins were extracted by cutting out each protein track into 1 piece followed by shotgun proteomic analyses. For western blot analyses, protein complexes were separated by SDS-PAGE on 4-20% pre-cast Bis-Tris gel (NuPAGE Life technology).

### Proteomic analysis

The mass spectrometry analyses were performed at Paris Sud Ouest PAPPSO proteomics core facility (http://papso.inra.fr). Briefly, proteins were digested for 6 h at 37°C with 4ng.μL^−1^ of modified trypsin (Promega) dissolved in 50 mM NH_4_CO_3_. Peptides were extracted with 2% trifluoroacetic acid (TFA) and 50% acetonitrile (ACN). Peptide extracts were dried and suspended in 15 μL of 0.05% TFA, 0.05% HCOOH, and 2% ACN. 4 μL of samples were then loaded on a NanoLC-Ultra system (Eksigent). Eluted peptides were analysed on-line with a QExactive mass spectrometer (Thermo Electron) using a nanoelectrospray interface. Peptide ions were analysed using Xcalibur 2.1. Peptide spectra from MS/MS analysis was identified with X!Tandem data base. A student test was performed to compare the specificity of proteins present in all experiment. Only candidate proteins with a t test inferior to 0.05 were retained.

### Multiple Sequence Alignments and Phylogenetic Tree Constructions

The complete archaeal and bacterial genome entries were retrieved from the EBI (European Bioinformatics Institute). The collection of helicases from super-families SF1 and SF2 were identified by RPS-Blast of Pfam domains DEAD (PF00270) or Helicase_C (PF00271) among a set of completely sequenced non-redundant genomes composed of 114 archaeal genomes and 1105 bacterial genomes. 17661 helicases were identified and further compared to each other using the BlastP program. Partitioning of the all-vs-all Blast results was achieved using the Markov Cluster Algorithm (71) and resulted in 84 clusters of >5 orthologous archaeal helicases. The archaeal sequences were collected and corresponded to occurrences with COG1202 for ASH-Ski2. We also collected an initial set of proteins using COG profile annotations (COG1096 for Csl4 and COG0689 for Rrp41) from complete archaeal genomes. Hidden Markov model (hmm) profiles were built for each family and used with hmmsearch (HMMER package, (72)) to identify homologs in our archaeal proteome sample. In addition, to identify potential unannotated genes and pseudogenes, we performed tblastn searches.

The archaeal phylogenetic tree was inferred from a concatenated dataset of 81 protein families obtained from COG annotation. If multiple copies occurred in a genome, all paralogs were removed. The sequence alignments for each family were created using the MUSCLE program (73) with the default parameters. We used the trimAl program (74) to remove spurious sequences and poorly aligned positions and to analyse the quality of the alignments according to gap numbers and residue conservation in the columns of the alignments. These parsed alignments were concatenated to produce a single alignment of 17827 residues. When a species did not have a record for a family, the missing sequence was replaced by gaps in the alignment. The maximum-likelihood tree was computed with PhyML (75) and the optimal combinations of parameters was selected using ProtTest3 program (76). The LG model of sequence evolution was used and the gamma-distributed substitution rate variation was approximated by eight discrete categories with shape parameter and proportion of invariant sites estimated from the data. The statistical branch support was inferred with the parametric bootstrap.

The archaeal Ski2-like helicase alignment was built by Mafft incorporating local pairwise refinement (L-INS-i) up to 2000 iteration (maxiterate 2000) (77) and trimmed with trimAL as described above. The archaeal ASH-Ski2 helicase tree was computed with the same approach as that of the species tree except that the gamma-distributed substitution rate variation was approximated by four discrete categories. Both trees (species and Ski2-like helicases) were arbitrarily rooted and were drawn with the online version of iTOL (78).

### Sucrose gradient fractionation of cell extracts

*P. abyssi* and *T. barophilus* cells were grown at 92°C and 85°C, respectively, in continuous culture in a 5L Gazlift bioreactor in MES medium under anaerobic conditions at pH 6.8 (79). Cells were maintained in exponential growth phase between 2 and 4 *10^8^ cells per mL. After cell culture harvesting at 8°C, Cells were pelleted by centrifugation (10,000g, 20min at 4°C) and washed two times with a sterile sea salt solution at 30g.L^−1^. Dry cell pellets were stored at −80°C. 200mg of cell pellet was re-suspended in 1/3 w/v of buffer TK buffer (20mM Tris-HCl, 100mM KCl, 10mM MgCl_2_, 1mM DTT) containing a cocktail of EDTA-free protease inhibitor (cOmpleteTM, Roche) or TK-EDTA buffer (TK buffer supplemented with 20mM EDTA). Whole-cell extracts were prepared by sonication (10*10 sec, 50% cycle, VibraCell-Bioblock scientific). Lysates were cleared at 21,000g for 30 min. ~10mg of whole-cell extract protein were layered on a linear 10-30 % sucrose gradient prepared in TK or TK-EDTA buffers and centrifuged for 5h at 35,000 rpm at 4°C in Beckman-Avanti XPN-80 SW41 rotor. It was not possible to determine the positions of 30S and 50S ribosomal subunits by classical A254 scanning with the ISCO UA-6 gradient fraction collector due to the inherent colorimetry of the *P. abyssi* and *T. barophilus* cellular extract.

For each gradient, 21 fractions of 500μL were collected. 250 μL of each fraction were precipitated with of 45μL of 100 % trichloacetic acid (TCA). After 30 min at −20°C, samples were centrifuged 30 min at 16,000g at 4°C. The protein pellets were washed with 500μL of glacial acetone (16,000g, 15min at 4°C), dried and re-suspended in 20μL of 20 mM Tris-HCL pH 8. After adding 4μL of 6X loading buffer (375mM Tris-HCL pH 6.8, 12% SDS, 30% β-mercaptoethanol, 0.6 % Bromophenol Blue, 36% glycerol), samples were heated at 95°C for 10 min before proceeding to SDS-PAGE.

### Relative quantification of ribosomal RNAs by Slot Blot

20 μL of each fraction collected from sucrose gradient, equally diluted beforehand to set the fraction with the highest nucleic acid content at a concentration of 2.5ng.μL^−1^, was added to 55 μL of denaturation buffer (2.2 M formaldehyde, 50% formamide, 0.5 mM EDTA, 10 mM MOPS, 4 mM NaCl) and incubated at 65°C for 5 min. Each 75 μL-sample was spotted on a nylon membrane (Amersham Hybond-XL; GE Healthcare) by vacuum filtration (PR648-HoeferTM Slot Blot). After UV-crosslinking, blots were hybridized overnight at 42°C with 5’-end P^32^-labelled oligonucleotides in Roti-Hybrid-Quick buffer (Roth) and washed three times at 42°C for 15min with SSC buffer (SSC 20X: 3M NaCl, 300mM sodium citrate, pH 7) containing 0.1% of SDS (5X, 1X and 0.1X, respectively). Sequences of antisense oligonucleotides used to detect *T. barophilus* 16S and 23S rRNAs are listed in **Table S4**. Radioactive signals were visualized on a phoshorImager (Typhoon Trio-Amersham-Bioscience) and quantified using MultiGauge software (Fujifilm).

## ACKNOWLEDGMENTS

We are indebted to the Paris Sud Ouest PAPPSO proteomics core facility (http://papso.inra.fr) for LC/MS analyses. We thank Y. Quentin for its expertise in taxonomic identification of archaeal Csl4 and Rrp41 members, L.Plassart and J. Caumes for technical help and P. Vitali and M. Kwapisz for helpful discussions.

## AUTHOR CONTRIBUTIONS

Conceived and designed the experiments: B.C.O., D.F., M.Bo., M.J., C.E., D.K.P. Conceived and designed the phylogenomic analyses: G.F., P.L.G.. Performed the biological experiments: D.K.P., C.E., M.Ba., Y.M., S.L., V.M. Performed the phylogenomic analyses: P.L.G. Analysed the data: D.K.P., B.C.O., D.F., M.Bo. Wrote the paper: B.C.O., D.F., M.Bo., D.K.P., C.E.. All authors read and approved the final manuscript.

