## APPENDIX Tables and Figures for "RNA processing machineries in Archaea: the 5’-3’ exoribonuclease aRNase J of the β-CASP family is engaged specifically with the helicase ASH-Ski2 and the 3’-5’ exoribonucleolytic RNA exosome machinery"

#### TABLE S1

| ACCESSION | GENE | NAME | M | AA | # NUC- | # NUC+ | DESCRIPTION |
|---|---|---|---|---|---|---|---|
| Q9UZ78 | PAB2428 | **TmcA** | 94 | 817 | 23 ± 9 | 27 ± 14 | tRNA(Met) cytidine acetyltransferase |
| Q9V1F2 | PAB0316 | **DnaG** | 50 | 447 | 32 ± 5 | 29 ± 4 | RNA exosome cap subunit |
| Q9V115 | PAB0423 | **RpoB** | 127 | 1117 | 11 ± 1 | 6 ± 1 | DNA-directed RNA polymerase subunit beta |
| Q9V1T4 | PAB2119 | ***Unch.*** | 29 | 258 | 7 ± 1 | 7 ± 1 | Uncharacterized methyl-transferase like |
| Q9V1Y9 | PAB2163 | **RPA41** | 43 | 375 | 7 ± 1 | 6 ± 1 | RPA41 subunit |
| Q9V2L4 | PAB2305 | **Nop5p** | 47 | 404 | 6 ± 3 | 7 ± 1 | Nop58p-like C/D snoRNP |
| Q9UYB6 | PAB1306 | **eIF-2B** | 31 | 276 | 5 ± 4 | 4.0 ±1.5 | Translation initiation factor subunit 2-like |
| Q9V2M1 | PAB2313 | **ASH-Ski2** | 97 | 855 | 5 ± 2 | 2 ± 1 | ATP-dependent RNA helicase |
| Q9V1T5 | PAB2120 | **Rpl3** | 40 | 361 | 4 ± 2 | 8 ± 3 | 50S ribosomal protein L3 |
| Q9UY07 | PAB1115 | **GraD-2** | 46 | 419 | 4.0 ± 0.5 | 5 ± 1 | Glucose-1-phosphate thymidylyltransferase |
| Q9UY85 | PAB1284 | **RecJ-like** | 83 | 740 | 4 ± 1 | 4 ± 1 | RecJ-like phosphoesterase |
| Q9V2L5 | PAB2306 | **FlpA** | 25 | 227 | 15.0 ± 0.5 | 9 ± 3 | Fibrillarin -rRNA/tRNA 2'-O-methyltransferase |
| Q9V119 | PAB0420 | **Rrp41** | 27 | 249 | 8 ± 2 | 2 ± 1 | RNA exosome catalytic core subunit |
| Q9V2F9 | PAB0071 | **AubA.** | 53 | 469 | 6 ± 1 | - | RNA-binding protein AU-1 |
| Q9V118 | PAB0421 | **Rrp42** | 29 | 274 | 5 ± 1 | 2.0 ± 1.5 | RNA exosome core subunit |
| Q9V120 | PAB0419 | **Rrp4** | 29 | 265 | 5 ± 1 | 2.0 ± 1.5 | RNA exosome cap subunit |
| Q9UZN6 | PAB1633 | **PINA** | 68 | 608 | 5 ± 3 | 7 ± 5 | PIN domain ATPase |
| Q9V113 | PAB0425 | **RpoA2** | 44 | 397 | 4 ± 2 | 1 ± 1 | DNA-directed RNA polymerase subunit A" |
| Q9V114 | PAB0424 | **RpoA1** | 103 | 907 | 4 ±1 | 2.0 ± 0.5 | DNA-directed RNA polymerase subunit A’ |

Most significant protein candidates of *Pab*-aRNase J-His detected by bottom-up proteomic techniques coupled with mass spectrometry. Spectra average numbers (#), from triplicate assays, rounded down for values below 0.5 and up for values above 0.5 are indicated. Standard error of the mean of three independent experiments is indicated for each value. Candidates with a maximal total number of spectra higher than 5 in either condition are considered. Candidates detected have a p-value lower than 0 (upper part) or 0.05 (lower part). Proteins accessions are from Uniprot data base (http://www.uniprot.org/uniprot). Nuc- and Nuc+ are for assay in absence and in presence of RNase/DNase treatment, respectively. Molecular weight in kDa (M), uncharacterized proteins (Unch.). Partners described to be involved in RNA synthesis, modification or decay and in translation are highlighted in light grey and, more specifically, the ASH-Ski2 helicase partner in darker grey.

ASH-Ski2 partners

#### TABLE S2A

| ACCESSION | GENE | NAME | M | AA | # NUC- | # NUC+ | DESCRIPTION |
|---|---|---|---|---|---|---|---|
| Q9UZ78 | PAB2428 | **TmcA** | 94 | 817 | 40 ± 2 | 40.5 ±1.0 | tRNA(Met) cytidine acetyltransferase |
| Q9V089 | PAB2390 | **Sun protein** | 52 | 450 | 37 ± 0 | 27.5 ±1.0 | Sun protein (Fmu protein) |
| Q9V1F2 | PAB0316 | **DnaG** | 50 | 447 | 16.5 ± 1 | 13 ±3 | RNA exosome cap subunit |
| Q9V2L4 | PAB2305 | **Nop5p** | 47 | 404 | 15.5 ± 1 | 13.0 ±1.5 | Nop58p-like C/D snoRNP |
| Q9V2L5 | PAB2306 | **FlpA** | 25 | 227 | 15.5 ± 6 | 11 ± 0 | Fibrillarin -rRNA/tRNA 2'-O-methyltransferase |
| Q9V192 | PAB0367 | **Eno-like** | 37 | 342 | 11.5 ± 1 | 11.5 ± 3.5 | Eno-like enolase related |
| Q9V076 | PAB1751 | **aRNase J** | 49 | 451 | 11 ± 7 | 8 ±0 | 5’-3’ Exoribonuclease |
| P62008 | PAB0460 | **Rpl7ae** | 13 | 123 | 11 ± 7 | 3.0 ±1.5 | 50S ribosomal protein L7Ae |
| Q9V115 | PAB0423 | **RpoB** | 127 | 1117 | 10.5 ± 2 | 6.5 ±3.5 | DNA-directed RNA polymerase subunit beta |
| Q9V1V5 | PAB2136 | **Rps5** | 26 | 236 | 8.5 ± 2.0 | 8.0 ±1.5 | 30S ribosomal protein S5 |
| Q9V1V6 | PAB2137 | **Rpl30** | 17 | 155 | 6.5 ± 2 | 4.5 ±2.0 | 50S ribosomal protein L30 |
| Q9UY85 | PAB1284 | **RecJ-like** | 83 | 740 | 6.5 ± 5.0 | - | RecJ-like phosphoesterase |
| G8ZHS0 | PAB2163 | **RPA41** | 40 | 358 | 5.5 ± 2.0 | 5 ± 0 | RPA41 subunit |
| Q9V236 | PAB0161 | **Unch.** | 23 | 204 | 5 ± 3 | 2 ± 0 | Uncharacterized protein |
| P61992 | PAB0361 | **Rps4** | 23 | 180 | 4.5 ± 3.5 | 3 ± 0 | 30S ribosomal protein S4 |
| Q9V1T8 | PAB2122 | **Rpl2** | 26 | 239 | 4.5 ± 2 | 2.5 ±1.0 | 50S ribosomal protein L2 |
| Q9UXX7 | PAB1136 | **Rnp3** | 24 | 212 | 4.5 ± 2 | 2.5 ± 1.0 | Ribonuclease P component 3 |
| Q9UWR8 | PAB1166 | **Rpl1** | 24 | 219 | 4.5 ± 1.0 | - | 50S ribosomal protein L1 |
| Q9V0G8 | PAB1813 | **Rps19** | 17 | 150 | 3.5 ± 2 | 2 ± 0 | 30S ribosomal protein S19 |
| Q9UZN6 | PAB1633 | **PINA** | 68 | 608 | 3 ± 0 | 4.5 ±1.0 | PIN domain ATPase |
| Q9V1N3 | PAB7094 | **AlbA** | 10 | 93 | 31.5 ± 10 | 7.5 ±2.0 | DNA/RNA-binding protein Alba |
| Q9UYV2 | PAB0931 | **CysS** | 56 | 477 | 12.5 ± 5 | - | Cysteine tRNA ligase |
| Q9V120 | PAB0419 | **Rrp4** | 29 | 265 | 11.5 ± 0 | - | Cap RNA exosome subunit Rrp4 |
| Q9V118 | PAB0421 | **Rrp42** | 29 | 274 | 9 ± 3 | - | RNA exosome core subunit |
| Q9V119 | PAB0420 | **Rrp41** | 27 | 249 | 7 ± 1.5 | - | RNA exosome catalytic core subunit |
| Q9V2F9 | PAB0071 | **AubA** | 53 | 469 | 7 ± 1.5 | - | RNA-binding protein AU-1 |
| Q9UZF3 | PAB1584 | **Unch.** | 19 | 164 | 5.5 ± 2 | - | Uncharacterized protein |
| Q9V1A5 | PAB05 | **TruB** | 38 | 334 | 5 ± 1.5 | 3 ± 0 | tRNA pseudo uridine synthetase |

Most significant protein partners of *Pab*-ASH-Ski2-His detected by bottom-up proteomic techniques coupled with mass spectrometry. See Table S1 for legends. Partners described to be involved in RNA synthesis, modification or decay and in translation are highlighted in light grey and, more specifically, the aRNase J partner in darker grey.

#### TABLE S2B

| ACCESSION | GENE | NAME | KDA | AA | # NUC- | # NUC+ | DESCRIPTION |
|---|---|---|---|---|---|---|---|
| Q9V115 | PAB0423 | **RpoB** | 127 | 1117 | 37.0 ± 1.5 | 25 ± 4 | DNA-directed RNA polymerase subunit beta |
| Q9UZ78 | PAB2428 | **TmcA** | 94 | 817 | 35.0 ± 1.5 | 35 ± 7 | tRNA(Met) cytidine acetyltransferase |
| Q9V192 | PAB0367 | **Eno-like** | 37 | 342 | 25 ± 4 | 21 ± 5.5 | Eno-like enolase related |
| G8ZHS0 | PAB2163 | **RPA41** | 43 | 375 | 23 ± 0 | 22 ± 0 | RPA41 subunit |
| Q9UZ86 | PAB2423 | **Rgy** | 140 | 1214 | 17 ± 7 | 14.0 ± 5.5 | Reverse gyrase |
| Q9V1Z1 | PAB2165 | **RPA32** | 31 | 272 | 16.5 ± 3.5 | 16 ± 4 | RPA32 subunit |
| Q9V133 | PAB2412 | **Unch.** | 48 | 415 | 15.0 ± 1.5 | 15.5 ± 5.0 | DUF530 |
| Q9V1F2 | PAB0316 | **DnaG** | 50 | 447 | 13 ± 1.5 | 11.5 ± 1.0 | RNA exosome cap subunit |
| Q9V2L4 | PAB2305 | **Nop5p** | 47 | 404 | 12.5 ± 1 | 13.5 ± 2.0 | Nop58p-like C/D snoRNP |
| Q9V114 | PAB0424 | **RpoA1** | 103 | 907 | 12 ± 0 | 6.5 ± 2.0 | DNA-directed RNA polymerase subunit A’ |
| Q9V2L5 | PAB2306 | **FlpA** | 25 | 227 | 11.5 ± 1.0 | 11 ± 1.5 | Fibrillarin-like rRNA/tRNA 2'-O-methyltransferase |
| Q9V076 | PAB1751 | **aRNase J** | 49 | 451 | 10.5 ± 2.0 | 8 ± 0 | 5’-3’ Exoribonuclease |
| Q9UY85 | PAB1284 | **RecJ-like** | 83 | 740 | 9.5 ± 2.0 | 9 ± 1.5 | RecJ-like phosphoesterase |
| P62008 | PAB0460 | **Rpl7ae** | 13 | 123 | 7.5 ± 1.0 | 3.5 ± 1.0 | 50S ribosomal protein L7Ae |
| Q9UZD0 | PAB0810 | **Predicted ATPase** | 61 | 548 | 6.5 ± 1.0 | 6 ± 1.5 | Predicted ATPase |
| G8ZJS6 | PAB0744 | **Lhr2** | 100 | 867 | 6.0 ± 1.5 | 6.5 ± 1.0 | Lhr-2 large helicase-related |
| Q9V089 | PAB2390 | **Sun protein** | 52 | 450 | 6 ± 0 | 5.0 ± 1.5 | Sun protein (Fmu protein) |
| Q9V1V5 | PAB2136 | **Rps5** | 26 | 236 | 5.5 ± 1.0 | 4.5 ± 1.0 | 30S ribosomal protein S5 |
| Q9V113 | PAB0425 | **RpoA2** | 44 | 397 | 5.5 ± 1.0 | 4 ± 0 | DNA-directed RNA polymerase subunit A" |
| Q9V1T5 | PAB2120 | **Rpl3** | 40 | 361 | 3.5 ± 1.0 | 4.0 ± 1.5 | 50S ribosomal protein L3 |
| Q9UYS8 | PAB1430 | **TopA** | 78 | 685 | 3.5 ± 1.0 | 3.5 ± 2.0 | DNA topoisomerase 1 |
| Q9V1N3 | PAB7094 | **AlbA** | 10 | 93 | 23.5 ± 3.5 | 9.5 ±1 | DNA/RNA- binding protein Alba |
| Q9V2F9 | PAB0071 | **AubA.** | 53 | 469 | 9.0 ± 1.5 | - | RNA-binding protein AU-1 |
| Q9V120 | PAB0419 | **Rrp4** | 29 | 265 | 8.5 ± 2.0 | - | RNA exosome cap subunit |
| Q9V118 | PAB0421 | **Rrp42** | 29 | 274 | 4 ± 3 | - | RNA exosome core subunit |

Most significant protein partners of *Pab*-His-ASH-Ski2 detected by bottom-up proteomic techniques coupled with mass spectrometry. See Table S1 for legends. Partners described to be involved in RNA synthesis, modification or decay and in translation are highlighted in light grey and, more specifically, the aRNase J partner in darker grey.

**TABLE S3**

| GENES | PROTEINS | N-His pET15b | C-His pET21b | pET11b |
|---|---|---|---|---|
| PAB1751 | aRNase J | √ | √ | √ |
| PAB2313 | ASH-Ski2 | √ | | √ |
| | ΔC-ASH-Ski2 | √ | | |
| | ΔN-ASH-Ski2 | √ | | |
| | ΔNC-ASH-Ski2 | √ | | |
| | ASH-Ski2-C124A | √ | | |
| | DomN | √ | | |
| | DomN C124A | √ | | |
| PAB0592 | Hel308 | √ | | |
| PAB2314 | Csl4 | √ | | √ |
| | ΔC-Csl4 | √ | | |
| | ΔN-Csl4 | √ | | |
| | Csl4-F121A | √ | | |
| | Csl4-F129A | √ | | |
| PAB0316 | DnaG | | √ | |
| PAB 0419 | Rrp4 | √ | | |
| PAB0420 | Rrp41 | √ | | |

pET vector constructions used to express recombinant proteins in *E.coli*.

**TABLE S4**

| | OLIGONUCLEOTIDES | SEQUENCES (5'-3') |
|---|---|---|
| **Reverse PCR** | For pET15b | CATATGGCTGCCGCGCGGC |
| | Rev pET15b / pET11b | GGATCCGGCTGCTAACAAAGCC |
| | For pET11b | CATATGTATATCTCCTTCTTAAAGTT |
| **Insert PCR** | For aRNaseJ pET11b | GGAGATATACATATGTGGGAGGAGATAAACATGATCA |
| | Rev aRNaseJ pET11b | TTAGCAGCCGGATCCTCATCCCTCCAATGAGCCAG |
| | For ASH-Ski2 pET15b | CGCGGCAGCCATATGCTATTCGTTATTCGCCCAGGGAGG |
| | Rev ASH-Ski2 pET15b | AGCCGGATCCTCGAGTTATGGTTTTCTTTCTCTCCTCACCTT |
| | For ASH-Ski2 pET11b | GGAGATATACATATGATGCTATTCGTTATTCGCCCAG |
| | Rev ASH-Ski2 pET11b | TTAGCAGCCGGATCCTTATGGTTTTCTTTCTCTCCTCA |
| | For ASH-Ski2ΔN-ter (194- 855) pET15b | CCGCGCGGCAGCCATATGACGATTGACGAGCTGGAT |
| | Rev ASH-Ski2ΔC-ter (1-573) pET15b | AGCCGGATCCTCGAGTTAAGAGGTTAGCAATTTCAAGGCCAC |
| | Rev DomN (1-193) ASH-Ski2 pET15b | TTAGCAGCCGGATCCTTA TCTCTCAACTTTAATACGCTTC |
| | For Hel308 pET15b | CGCGGCAGCCATATGATGAAAGTTGGAGAGCTAAACG |
| | RevHel308 pET15b | TTAGCAGCCGGATCCTTATGGGTTCAGGAAATAGTCC |
| | For Csl4 pET15b / pET11b | CGCGGCAGCCATATGTTGGAGGAAGGTGAGGAGAG |
| | Rev Csl4 pET15b | TTAGCAGCCGGATCC TCATAGCTTCACCTTCCTGTA |
| | For Csl4 pET11b | GGAGATATACATATGTTGGAGGAAGGTGAGGAGAG |
| | For Csl4ΔN-ter pET15b | AGGATCCTTTGTCGTCAACTGAATC |
| | Rev Csl4ΔC-ter pET15b | CCTCCAATACCTAAGAAAGGGGA |
| | For DnaG pET21b | GGAGATATACATATGATGAAAAGAAAGAGGGCGATAAT |
| | Rev DnaG pET21b | GTGGTGCTCGAGTGCCTCGGCGAAGGTTATTACCTT |
| | For Rrp4 pET15b | CGCGGCAGCCATATG ATGAAGAGGATTTTTGTTCAAAAT |
| | Rev Rrp4 pET15b | TTAGCAGCCGGATCC TTAAGCCCCTCTGGCTTCTC |
| | For Rrp41 pET15b | CGCGGCAGCCATATGATGATGGAGAAGCCAGAGGG |
| | Rev Rrp41 pET15b | TTAGCAGCCGGATCCTCACTCACTTCCCTCAACCT |
| **Directed mutagenesis** | For aRNase J Δ1 pET15b | GATTAGAACCCTGTCGAGCCTTATTC |
| | Rev aRNase J Δ1 pET15b | GACCTTAGGAAGCTCGGTGCAATAC |
| | For ASH-Ski2 C124A | GAGAGTACATAGCCGAGAGATGTG |
| | Rev ASH-Ski2 C124A | CACATCTCTCGGCTATGTACTCTC |
| | For Csl4 F121A | GTTAAGGACGGCGCCGTTGAGGATTTAAGGAA |
| | Rev Csl4 F121A | TAAATCCTCAACGGCGCCGTCCTTAACTTGAG |
| **Slot Blot** | *Tba* 16S rRNA | CTCCACCCCTTGTAGTGCTC |
| | *Tba* 23S rRNA | CCACGGCTGACGAACATTGCC |

List of oligonucleotides used in this study

### Figure S1

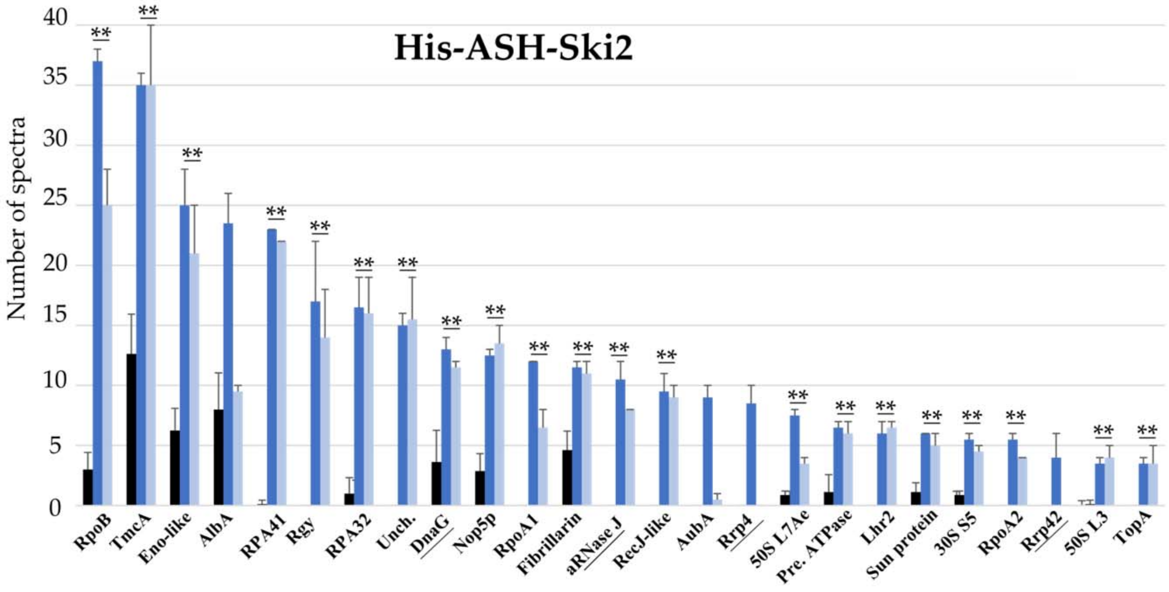


**FIG. S1** ASH-Ski2 partners from pull down assays using the recombinant *Pab*-(His)-ASH Ski2 protein as bait and *P. abyssi* cellular extract (see also Appendix Tabl. S2B). Legends as for Fig. 1.

### Figure S2

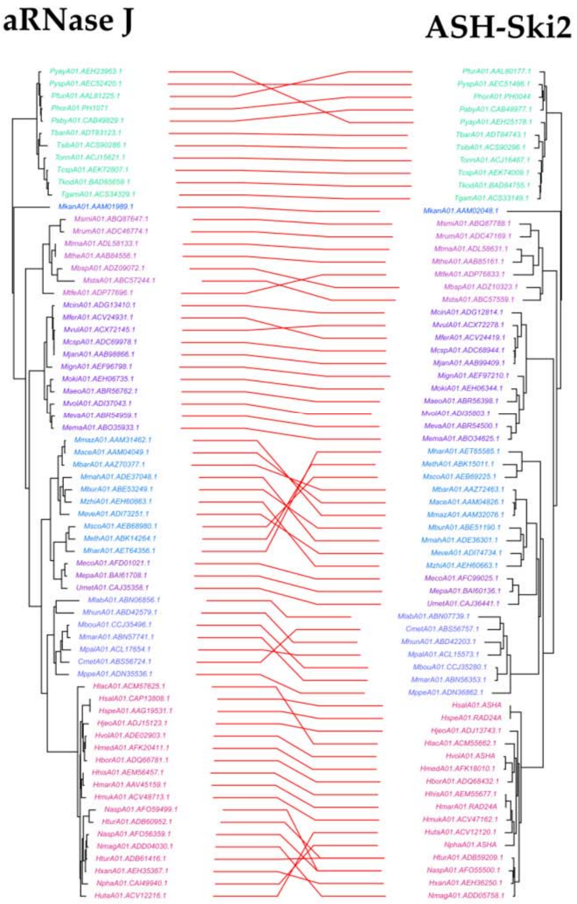


**FIG. S2** The aRNase J and ASH-Ski2 phylogenetic trees are congruent. Tips from the same genome are linked by a red line. Colour code for taxonomic order as in Fig. 3

### Figure S3

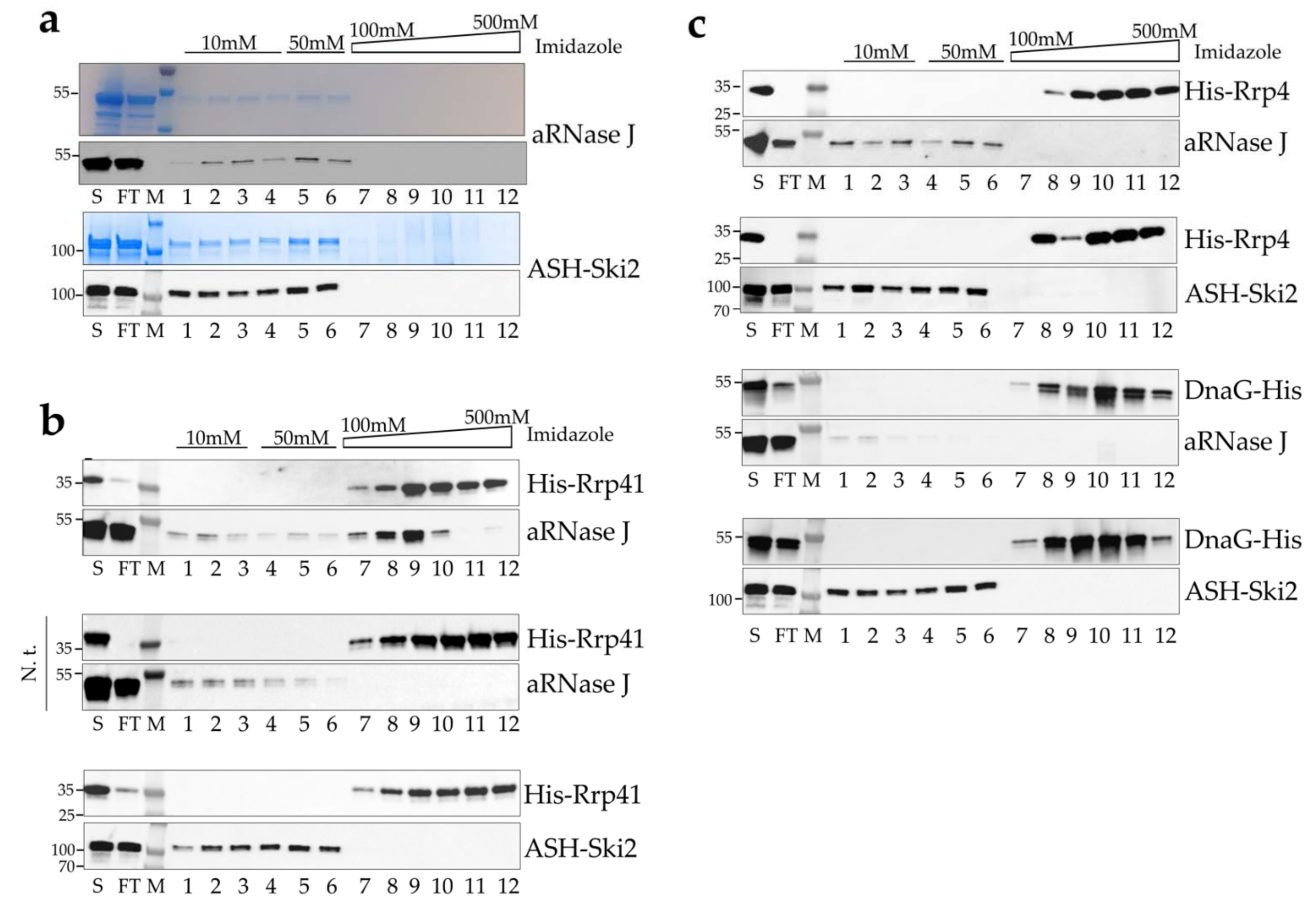


**Fig. S3** *In vitro* co-purification assays. **a** Affinity purification controls. Untagged *Pab*-aRNase J (top) and *Pab*-ASH-Ski2 (bottom) are not intrinsically retained on nickel column matrices. **b** Co-purification assays in which *Pab*-(His)$_6$-Rrp41 (bait) was challenged with *Pab*-aRNase J (prey) (left panel) or *Pab*-ASH-Ski2 (prey) (right panel), respectively. **c** Co-purification assays in which *Pab*-(His)$_6$-Rrp4 (bait) (left panel) or *Pab*-(His)$_6$-DnaG (bait) (right panel) was challenged with *Pab*-aRNase J (prey) or *Pab*-ASH-Ski2 (prey), respectively. Legend as in Fig. 5

### Figure S4

## a

Csl4 S1 domain motifs

co-distributed with aRNase J

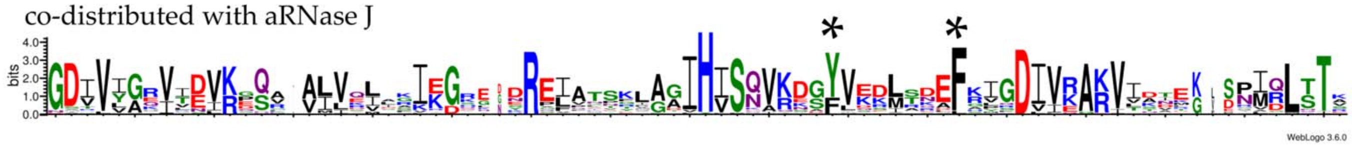


not distributed with aRNase J

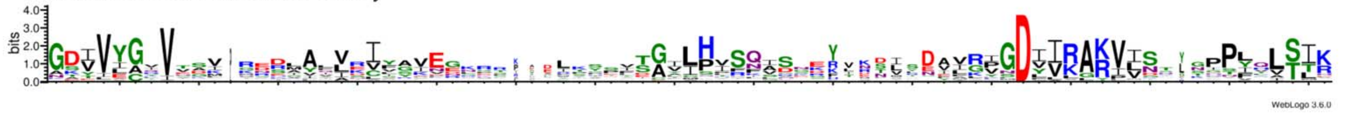


## b

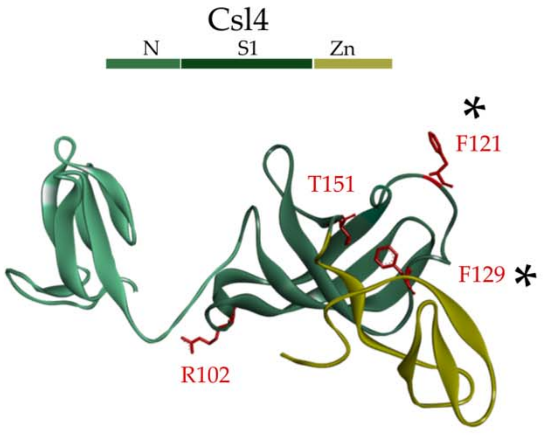


**FIG. S4** Csl4 structural domains. a Weblogo sequences derived from multiple alignments of Csl4 sequences co-distributed or not with aRNase J (referred to distribution of aRNase J & Csl4 displayed on Fig. 3). Several residues are specifically conserved in Csl4 sequences co-distributed with aRNase J. b The structural model for *P. abyssi* Csl4 is built using Phyre2 software. The solvent-exposed phenylalanine residues F121 and F129 are indicated by stars.

### Figure S5

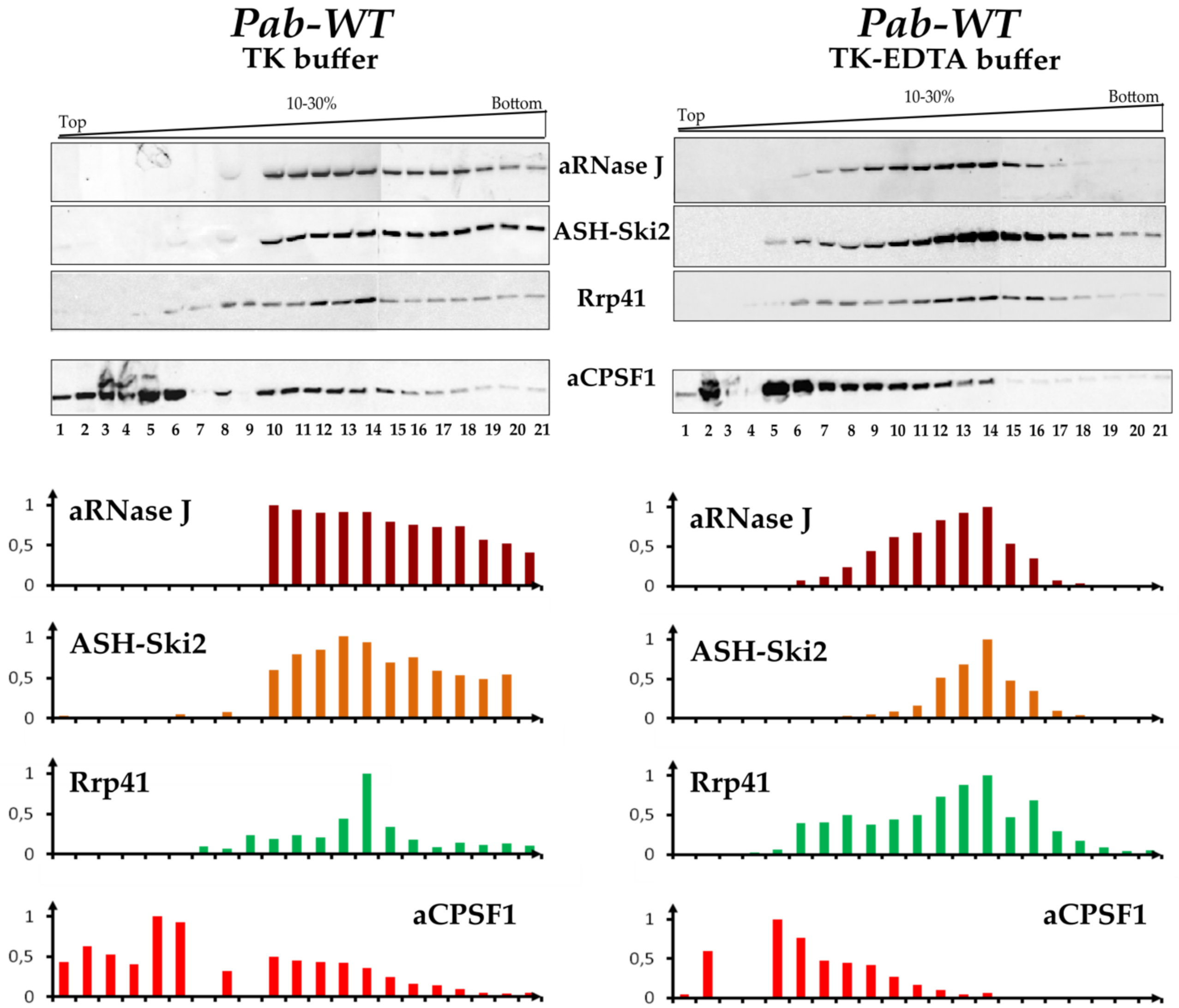


**FIG. S5** Sedimentation profiles of *Pab*-aRNase J, *Pab* -ASH-Ski2 and *Pab* -Rrp41 from WT strain *P. abyssi* cell extract performed in TK buffer in **a** and TK buffer supplemented with 20mM EDTA in **b**, as for Fig. 7. In here, the endogenous *Pab*-aCPSF1 β-CASP endo-RNase is also monitored as control.

### Figure S6

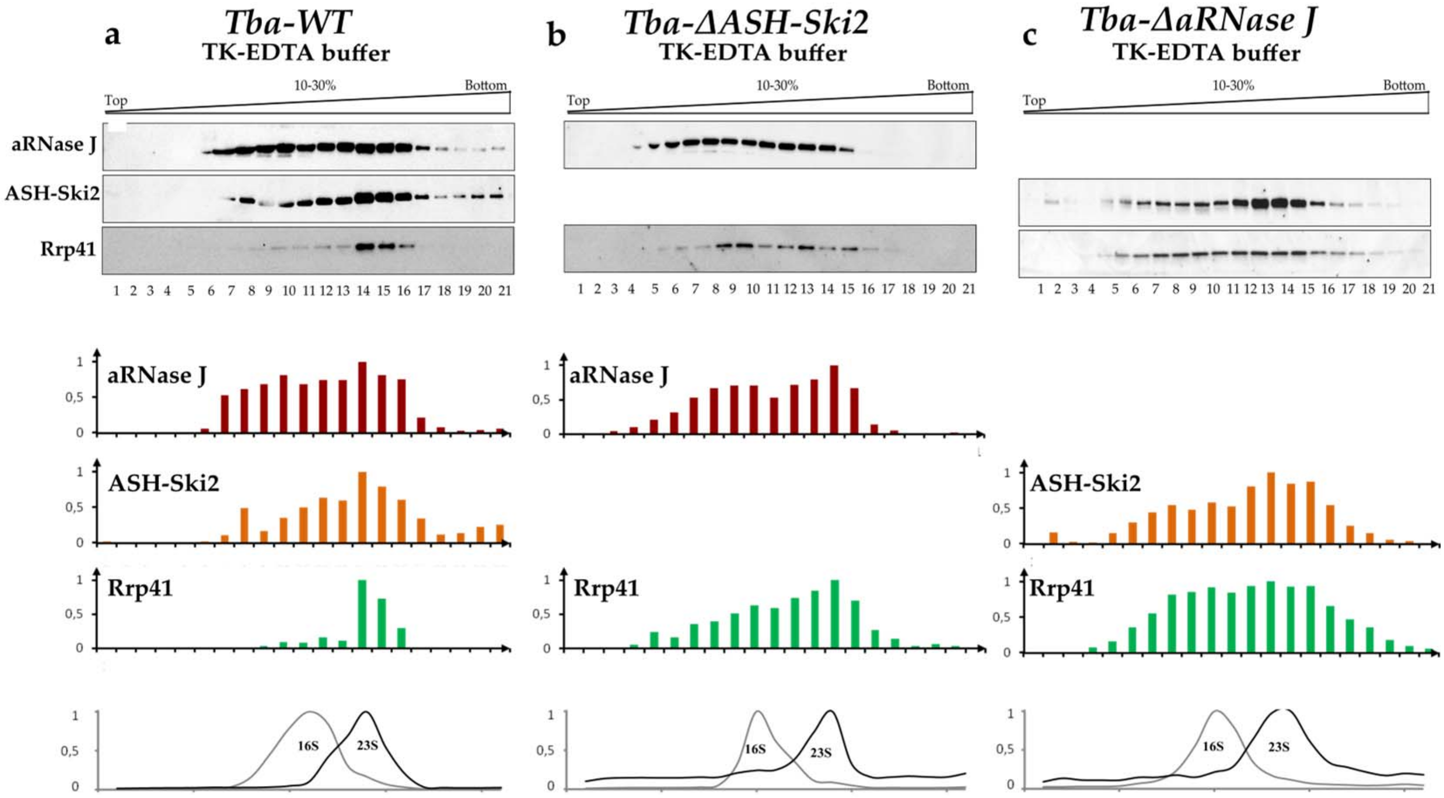


**FIG. S6** Sedimentation profiles of *Tba*-aRNase J, *Tba*-ASH-Ski2 and *Tba*-Rrp41 from the *Tba*-WT in **a**, *Tba*-ΔASH-Ski2 in **b**, and *Tba*-ΔaRNase J in **c**, strain cell extracts prepared in TK-EDTA buffer. Legend as for Fig. 6.
